# Single-cell transcriptomic profiling of kidney fibrosis identifies a novel specific fibroblast marker and putative disease target

**DOI:** 10.1101/2022.09.13.507855

**Authors:** Valeria Rudman-Melnick, Mike Adam, Kaitlynn Stowers, Andrew Potter, Qing Ma, Saagar M. Chokshi, Davy Vanhoutte, Iñigo Valiente-Alandi, Diana M. Lindquist, Michelle L. Nieman, J. Matthew Kofron, S. Steven Potter, Prasad Devarajan

**Affiliations:** Division of Nephrology and Hypertension, Cincinnati Children’s Hospital Medical Center, Cincinnati, OH, USA; Division Developmental Biology, Cincinnati Children’s Hospital Medical Center, Cincinnati, OH, USA; Division of Molecular Cardiovascular Biology, Cincinnati Children’s Hospital Medical Center, Cincinnati, OH, USA; Department of Pediatrics, University of Cincinnati, OH, USA; Cytokinetics, San Francisco, CA, USA; Department of Radiology, University of Cincinnati, OH, USA; Department of Radiology and Medical Imaging, Cincinnati Children’s Hospital Medical Center, Cincinnati, OH, USA; Department of Pharmacology and Systems Physiology, University of Cincinnati, OH, USA

**Author notes:** Corresponding author: Prasad Devarajan, 3333 Burnet Avenue, Cincinnati, OH 45229-3039, USA.

**Keywords:** chronic kidney disease, single cell, kidney fibroblast, kidney tubular injury, AHNAK

## Abstract

**Background:** Persistent kidney fibroblast activation and tubular epithelial cell (TEC) injury are key contributors to CKD. However, transcriptional and cellular identities of advanced kidney disease, along with renal fibroblast specific markers and molecular targets contributing to persistent tubular injury, remain elusive.

**Methods:** We performed single-cell RNA sequencing with two clinically relevant murine kidney fibrosis models. Day 28 post-injury was chosen to ensure advanced fibrotic disease. Identified gene expression signatures were validated using multiple quantitative molecular analyses.

**Results:** We revealed comprehensive single cell transcriptomic profiles of two independent kidney fibrosis models compared to normal control. Both models exhibited key CKD characteristics including renal blood flow decline, inflammatory expansion and proximal tubular loss. We identified novel populations including “secretory”, “migratory” and “contractile” activated fibroblasts, specifically labelled by newly identified fibroblast-specific *Gucy1a3* expression. Fibrotic kidneys elicited elevated embryonic and pro-fibrotic signaling, including separate “Embryonic” and “Pro-fibrotic” TEC clusters. Also, fibrosis caused enhanced cell-to-cell crosstalk, particularly between activated fibroblasts and pro-fibrotic TECs. Analysis of factors mediating mesenchymal phenotype in the injured epithelium identified persistent elevation of Ahnak, previously reported in AKI, in both CKD models. AHNAK knockdown in primary human renal proximal tubular epithelial cells induced a pro-fibrotic phenotype and exacerbated TGFβ response via p38, p42/44, pAKT, BMP and MMP signaling.

**Conclusions:** Our study comprehensively examined kidney fibrosis in two independent models at the singe-cell resolution, providing a valuable resource for the field. Moreover, we newly identified *Gucy1a3 as a* kidney activated fibroblast specific marker and validated AHNAK as a putative disease target.

**Significance Statement:** Mechanistic understanding of kidney fibrosis is principal for mechanistic understanding and developing targeted strategies against CKD. However, specific markers and molecular targets of key effector cells - activated kidney fibroblasts and injured tubular epithelial cells - remain elusive. Here, we created comprehensive single cell transcriptomic profiles of two clinically relevant kidney fibrosis models. We revealed “secretory”, “contractile” and “migratory” fibroblasts and identified Gucy1a3 as a novel marker selectively labelling all three populations. We revealed that kidney fibrosis elicited remarkable epithelial-to-stromal crosstalk and pro-fibrotic signaling in the tubular cells. Moreover, we mechanistically validated AHNAK as a putative novel kidney injury target in a primary human *in vitro* model of epithelial-to-mesenchymal transition. Our findings advance understanding of and targeted intervention in fibrotic kidney disease.

## Introduction

Fibrosis is a key underlying process in CKD, resulting in a progressive functional decline with high prevalence, morbidity and mortality^1–4^. While early fibrotic response is essential for injury recovery^5^, excessive extracellular matrix (ECM) production leads to renal parenchymal fibrotic remodeling^6^. Since existing therapeutic options remain merely supportive^7^ and advanced CKD might result in ESKD requiring lifelong dialysis or transplant^8^, mechanistic understanding of kidney fibrosis is paramount.

Stromal cells, particularly fibroblasts, are key contributors to kidney fibrosis^9^. Persistent injury triggers fibroblast activation and appearance of myofibroblasts, depositing excessive ECM and thereby causing fibrotic remodeling^10–12^. However, existing approaches to ECM producing renal cell populations targeting remain controversial, due to the nonspecific expression patterns of currently used markers. *Acta2*, encoding alpha 2 smooth muscle actin (αSma)^13^ is used as a main myofibroblast marker while also expressed in off-target renal cell populations including injured tubular epithelial cells (TECs)^14–19^. ECM components including Collagen1a1 (Col1a1) and Vimentin (Vim)^20^ also do not selectively label activated fibroblasts^21–24^. Thus, novel approaches allowing for specific renal activated fibroblast targeting are needed. Moreover, animal studies revealed remarkable transcriptional heterogeneity of renal stroma^25,26^ which represents another challenge for successful targeting. Thus, single cell resolution studies are essential to dissect the specific gene expression signatures of baseline and activated kidney fibroblasts.

Progressive mesenchymal changes in TECs represent another crucial process contributing to CKD^27^. Our recent single cell RNA sequencing (scRNA-seq) study showed remarkable induction of a pro-fibrotic phenotype in the TECs along with dramatically increased epithelial-to-stromal crosstalk after AKI^28–30^. Another single-nuclear transcriptomic study identified that failed proximal tubular recovery elicits pro-fibrotic secretory phenotype in the long-term kidney injury^1^. However, the molecular factors which orchestrate the persistent epithelial fibrogenic phenotype in CKD remain elusive.

Here, we used scRNA-seq to create comprehensive transcriptomic profiles of two clinically relevant murine kidney fibrosis models. We reproducibly observed several novel injury induced kidney cell clusters, including three distinctive stromal populations with “secretory”, “contractile” and “migratory” transcriptional enrichments. We newly identified and validated Gucy1a3 as a specific marker of renal quiescent and activated fibroblasts. Renal fibrotic injuries elicited a robust nephrogenic signature, enhanced epithelial-to-stromal interactions and persistent mesenchymal changes in the TECs. Moreover, we identified AHNAK as a putative novel kidney injury target in the primary human *in vitro* model of epithelial-to-mesenchymal transition (EMT).

## Methods

### Animals

The Institutional Care and Use Committee (IACUC) of Cincinnati Children’s Hospital Medical Center reviewed and approved all animal procedures used in this study. UIR was induced via atraumatic left renal pedicle clamping for 30 minutes at 37°C and UUO was induced via irreversible left ureter ligation in 10 weeks old male C57Bl/6 mice (n=3 for 10x Chromium scRNA-seq, 1 for Drop-seq per procedure). The kidneys were harvested at Day 28 post-injury. Normal mice of the same age, strain and sex were used as controls and harvested at the identical time (14 weeks old, n=5). Validations were performed on additional sets of animals treated with the identical UIR and UUO and on the leftover tissue from the scRNA-seq suspensions.

### scRNA-seq procedure

UIR, UUO and control mice were intraperitoneally injected with 100 μL heparin (100 U/mL), anesthetized with isoflurane chamber and euthanized via exsanguination. The animals were perfused with ice-cold PBS via the aorta, the left kidneys were decapsulated and minced finely with a sterile scissors until tissue was homogenous and clumps were broken down. 65 mg of the minced tissue was placed in 2 ml of ice-cold digestion buffer (10 mg/mL of cold active protease from *Bacillus licheniformis* (Sigma, P5380), 5mM CaCl2 and 125U/ml of DNAse 1 in PBS)^31^. The digest mix was incubated on ice and triturated with 1ml pipet tip until the clumps were no longer observed (15 sec every 2 min). Then, the cell suspensions were transferred to 15ml conical tubes and 3ml of ice-cold 10% FBS/PBS was added to inhibit the enzyme activity. The cell suspensions were filtered through 30μM Miltenyi filter rinsed with 4ml of ice-cold 10% FBS/DPBS to ensure the maximal cell yield and pelleted by centrifugation at 300 g for 5 minutes at 4°C. The supernatant was discarded, and the cellular pellet re-suspended in 1ml of ice-cold 0.01% BSA/PBS for Drop-seq and 10% FBS/DPBS for 10x Chromium at 100,000 cells/mL and to 1,000,000 cells/ml, respectively. The remaining minced tissue was snap-frozen in liquid nitrogen for molecular analysis. The contralateral kidneys were fixed with 4% paraformaldehyde (PFA) in PBS overnight (O/N) at 4°C and paraffin embedded for histological assessment. Independent cohorts of identical UIR, UUO and naïve mice (n=4-6 per group) were harvested at the same time point for validation.

### scRNA-seq data analysis

Libraries were sequenced using an Illumina Novaseq 6000. The raw fastqs were processed using CellRanger v3.1.0 (10X Genomics) aligning reads to the mm10 genome to generate a feature count matrix. Cell-type clusters and markers genes were identified using the R v4.1.1 library Seurat v3.2.3^32^. Initial cell filtering selected cells that expressed >500 genes. Genes included in the analysis were expressed in a minimum of three cells. Only one read per cell was needed for a gene to be counted as expressed per cell. Cells containing high percentages of mitochondrial, >30%, and hemoglobin genes, >2.5% were filtered out. The resulting gene expression matrix was normalized to 10,000 molecules per cell and log transformed. To minimize batch effect all eleven 10X runs, data sets were integrated using anchor genes. Integration features were found using the FindIntegrationFeatures function and integration anchors were found from these features using the FindIntegrationAnchors function. These anchors were used to integrate the data sets through the IntegrateData function. The final total number of 34875 cells was analyzed (Control: 9753, UIR: 13951, UUO: 11171). All clustering was unsupervised, without driver genes. The influence of the number of unique molecular identifiers was minimized by regression within the ScaleData function. Genes with the highest variability among cells were used for principal components analysis. Cell clusters were determined by the Louvain algorithm by calculating k-nearest neighbors and constructing a shared nearest neighbor graph, with a resolution set at 0.5. Dimension reduction was performed using UMAP (Uniform Manifold Approximation and Projection) using the first thirty principal components. Marker genes were determined for each cluster using the Wilcoxon Rank Sum test within the FindAllMarkers function using genes expressed in a minimum of 25% of cells and fold change threshold of 1.3. The trajectory analyses were performed on the populations designated in each graph using Monocle2^33^. Putative signaling interactions between proximal tubules, injured proximal tubules, stromal cells, and mixed-identity cells were assessed. Potential receptor-ligand interactions were found by pairing a cell-type expressing a ligand with a cell-type expressing its receptor pair. A receptor or ligand was considered expressed in a cell-type having an average expression of >0.24. Receptor-ligand pairs were determined using the curated receptor-ligand database by the RIKEN FANTOM5 project^34^. Receptor-ligand pairings for each cell type were visualized by a chord diagram using the R package circlize^35^. GO analysis was performed in the ToppGene Suite with 0.05 p value cutoff. The gene clusters enriched in renal cell populations were generated using ToppCluster with Bonferroni correction and 0.05 p value cutoff. The ToppCluster graph was generated with Fruchterman-Reingold graph layout algorithm, showing individual genes associated with biological processes enriched in the renal cell population of interest.

### Renal blood flow procedure (RBF)

The independent cohort of UIR, UUO and control mice was subjected to invasive hemodynamics. The mice were anesthetized with ketamine and inaction, femoral artery was cannulated for blood pressure monitoring and femoral vein was used for bolus infusion of medicines, including phenylephrine (PE) used for vasoconstriction, sodium nitroprusside (SNP) used for vasodilation, and dobutamine (DOB) for cardiac stress test. These medicines were administered to examine RBF alterations in the fibrotic and normal kidneys after altering the vascular tone and cardiac activity, as well as to measure basal cardiovascular parameters in the injured and control groups. To access the left kidney, a flank incision was made and the kidney was retracted to expose the renal artery. Then, an ultrasonic flow probe (Transonic systems) was placed around the left renal artery and positioned to obtain maximal blood flow. Mice were given 3-5 min between each dose to ensure the return to the baseline. 4% BSA at 0.15ul/min/gBW was used as maintenance infusion.

### Magnetic resonance imaging (MRI)

Mice were scanned using a horizontal 7T Biospec MRI system (Bruker, Billerica, MA) with a home-built 35 mm diameter quadrature volume transmit/receive coil. Mice were anesthetized with isoflurane and kept warm with circulating air, which was controlled by a temperature and respiration rate monitor (Small Animal Instruments, Inc., NY). Respiration was maintained around 100 breaths/minute. The mouse kidney was positioned at magnet and coil isocenter. Axial images were acquired using a fast spin echo sequence with a repetition time of 2500 ms, echo time of 40.2 ms, echo train length of 16, 4 averages, 32 mm x 32 mm field of view, and an acquisition matrix of 200 x 200.

### Primary Human RPTEC Culture, Transfection and TGFβ Treatment

Passage 3-4 RPTECs (CC2553) were cultured in the renal epithelial cell medium 2 (C-26130) supplemented with 5% fetal calf serum (FCS), 10ng/ml EGF, 5μg/ml human recombinant insulin, 0.5μg/ml epinephrine, 36ng/ml hydrocortisone, 5μg/ml recombinant human transferrin, 5pg/ml triiodo-L-thyronine, 30μg/ml Gentamycin and 15ng/ml Amphotericin. Once the cells reached near 70% confluency, transfection with ON-TARGETplus SMARTpool siRNAs targeting AHNAK or scramble non-targeting pool was performed in antibiotic-free conditions with DharmaFECT transfection reagent 1. 72 hours post-transfection, 5ng/ml TGFβ (R&D systems, 240-B-002) was given, and the cells were harvested at 1, 48 or 72 hours post-treatment. Sterile 4mM HCl with 1 mg/ml endotoxin-free BSA was used as Vehicle; primary human RPTECs cultured in identical conditions were used as controls. Effects of AHNAK knockdown on EMT in primary human RPTECs were validated in 5 independent experiments.

### Mass migration assay

Primary human RPTECs were cultured in 24 well plates (Ibidi, 82406), transfected with siScramble or siAHNAK as described above and placed in antibiotic-free starvation medium (0.5% FCS) 48 hours post-transfection for 24 hours. Then, scratch was performed with 200μl pipette tip, RPTECs were washed twice with pre-warmed sterile PBS, treated with Vehicle or 5ng/ml TGFβ in fresh antibiotic-free starvation medium and imaged for 24 hours on the Nikon Ti2 wide-field microscope. Obtained images were analyzed with ImageJ “Straight” line tool, and scratch closure coefficient (%) was determined at 18014.24, 40514.03, 63013.77, 85513.53 sec post-scratch via dividing the scratch size at a given time-point to the scratch size at 00.00 sec time point. Three coefficients obtained from each individual well were averaged, and then 4 wells per treatment group were averaged. The observation was reproduced in two independent runs (n=4 wells per group).

### Real-time quantitative PCR (RT-qPCR)

Total RNA was isolated from homogenized control, UIR and UUO whole kidney lysates (n=4-6 per group) with RNA Stat-60 extraction reagent (Amsbio, CS-111) and purified using the GeneJET RNA purification kit (ThermoFisher Scientific, KO732). Total RNA and protein were simultaneously isolated from RPTECs using Ambion PARIS kit (AM1921, ThermoFisher Scientific). cDNA was synthesized with the iScript Reverse Transcription Supermix (Bio-Rad, 1708841). qPCR was performed with TaqMan universal PCR master mix (Thermo Fisher Scientific, 4304437) on the Applied Biosystems Quant Studio 3 system. The reported Ct values are the mean of two cDNA sample replicates. The target gene Ct values were normalized to the eukaryotic 18S rRNA endogenous control and presented as the fold change.

### Chromogenic in situ hybridization (CISH)

CISH riboprobes for *Spp1* and *Myh9* were generated from gDNA or cDNA with PCR and labeled with digoxigenin (DIG) using DIG RNA labeling mix (Roche, 11277073910) by *in vitro* transcription. PFA fixed paraffin embedded (PFPE) 6 μm kidney sections were subjected to the CISH protocol as previously described^30,36^. Deparaffinated and dehydrated sections were incubated with Proteinase K (1ug/ml), fixed in 4% PFA, acetylated in 0.25% acetic anhydride and hybridized for 14-16 hours at 70°C with antisense or negative control sense riboprobes in RNase-free conditions. Then, sections underwent a series of saline sodium citrate buffer washes and were incubated with anti-DIG-AP fab fragments (Roche, 11093274910, 1:1000) O/N at 4 °C, followed by a series of post-hybridization washes and color development with BM Purple (Roche, 11442074001). For IF co-labeling, the protocol was modified by adding the primary antibody recognizing the target (Krt8, rabbit, Abcam, EP1628Y, 1/100) to the anti-DIG-AP antibody mixture. Next day, the sections were incubated with fluorescent secondary antibodies for 1 hour at room temperature (RT) in darkness, followed by the post-hybridization washes, color development, DAPI (Thermo Fisher Scientific, 62248, 1:1000) treatment for nuclei labeling and mounting with Vectashield antifade mounting medium (Vector Laboratories, H-1000). Secondary antibodies only application was used for the negative control. Imaging was performed on a Nikon A1R HD confocal TiE microscope using the galvanometric scanner. Transmitted images of chromogenic substrate are shown in gray-scale images.

### Single molecule fluorescent in situ hybridization (smFISH), using RNAscope

RNAscope probes and Multiplex Fluorescent v2 assay (323100) were purchased from Advanced Cell Diagnostics, Icn (ACD). Freshly sectioned PFPE 6 μm kidney sections underwent deparaffinization, dehydration, endogenous peroxidase quenching, heat-induced target retrieval and protease digestion, followed by incubation with up to four target riboprobes for 2 hours at 40°C. All the aforementioned steps were performed in RNase-free conditions. Next, tyramide signal amplification and conjugation to an Opal dye (PerkinElmer) were performed according to the manufacturer’s protocol. The sections were treated with DAPI and mounted with Vectashield antifade mounting medium. The controls were treated with negative control riboprobe and signal amplification reagents alongside the experimental sections. Images were obtained on 60x water immersion (WI) objective at Nyquist resolution on the Nikon Ti-E A1R HD confocal with the resonant scanner and processed with NIS-Elements AR 5.2.00 artificial intelligence denoise algorithm^30^. All images within an experimental group were obtained with the same optical configurations.

### Western Blotting

Total protein was extracted from homogenized whole kidney lysates and RPTECs using mammalian protein extraction reagent (MPER, ThermoFisher Scientific, 78501) and Ambion PARIS kit, respectively. Extraction buffers were supplemented with protease (ThermoFisher Scientific, 78430) and phosphatase (Sigma-Aldrich, P5726, P0044) inhibitors. 10-15 μg of protein was separated via PAGE, transferred to PVDF membrane, blocked with 5% non-fat milk and incubated with the target recognizing primary antibodies O/N at 4°C. Then, the membranes were washed with Tris buffered saline 0.02% Tween (TBST) and incubated with the secondary HRP-conjugated antibodies for 1 hour at RT. For phosphorylated target detection, PhosphoBLOCKER reagent (Cell Biolabs, AKR-103) was used during membrane blocking, primary and secondary antibody incubation. The target protein levels were normalized to the endogenous control detected with goat anti-Gapdh (AF5718, 1:200) or mouse anti-Gapdh (MAB374, 1:5000) antibodies. The signal was visualized using SignalFire ECL reagent (Cell Signaling, #6883). Following primary antibodies were used: rabbit anti-Ahnak (16637-1-AP, 1:250), rabbit anti-Sox4 (C15310129, 1:1000), rat anti-Cd24 (ab64064, 1:100), rabbit anti-Endostatin (ab207162, 1:1000), goat anti-Cd45 (AF114, 1:2000), rabbit anti-Fn (15613-1-AP, 1:1000), mouse anti-αSma (A5228, 1:1000), rabbit anti-Vim (ab45939, 1:1500), mouse anti-Ncad (610921, 1:1000), rabbit anti-Ecad (20874-1-AP, 1:5000), mouse anti-Ggt1 (ab55138, 1:750), rabbit anti-P-p38/p38 (4511S, 1:1000), mouse anti-P-Akt (4051S, 1:1000), rabbit anti-P-p44/42 (4376S, 1:1000), rabbit anti-P-Smad2 (3108S, 1:600), rabbit anti-P-Smad3 (NBP1-77836, 1:500), rabbit anti-p38 (9212S, 1:1000), rabbit antip44/42 (9102S, 1:1000), goat anti-Smad2/3 (AF3797, 1:400), goat anti-Bmp7 (sc9305, 1:100).

### Immunofluorescence (IF), immunohistochemistry (IHC), immunocytochemistry (ICC)

IF and ICC staining of fresh PFPE 6 μm kidney sections was performed via deparaffinization, heat induced citrate epitope retrieval, permeabilization in 0.06% Triton-X-100, blocking with 5% serum and O/N incubation with primary antibodies at 4°C. For IF, sections were then incubated with secondary fluorescent antibodies, stained with DAPI, mounted with Vectashield antifade mounting medium and imaged on 60x WI objective at Nyquist resolution on the Nikon Ti-E A1R HD confocal with the resonant scanner. For IHC, primary antibody incubation was followed by signal detection with the ImmPRESS reagent kit with DAB substrate^37^ and imaging on the Nikon Ti2 wide-field microscope. Following primary antibodies were used for IF: rabbit anti-Gucy1a3 (12605-1-AP, 1:50), guinea pig anti-Nphs (GP-N2, 1:50), mouse anti-αSma (A5228, 1:100), goat anti-Ecad (AF748, 1:20), mouse anti-Aqp1 (sc25287, 1:50), rabbit anti-Ahnak (16637-1-AP, 1:100). Following primary antibodies were used for ICC: rabbit anti-Sox4 (AB5803, 1:100), rabbit anti-Ahnak (16637-1-AP, 1:200), rabbit anti-Col18a1 (HPA011025, 1:50).

ICC was performed on primary human RPTECs cultured on Ibidi 12-well removable chamber slides (81201) and fixed with 4% PFA for 15 min at RT. RPTECs were permeabilized with 0.1% Triton-X-100, blocked with 2% BSA for 1 hour at RT, incubated with primary antibodies (mouse anti-AHNAK, ab68556, 1:250 and rabbit anti-ZO1, 21773-1-AP, 1:1000) at 4°C O/N and then with secondary fluorescent antibodies for 1 hour at RT, stained with DAPI and mounted with Vectashield antifade mounting medium. Images were obtained on 60x WI objective at Nyquist resolution on the Nikon Ti-E A1R HD confocal with the resonant scanner.

### Data availability

scRNA-seq data were deposited at the Gene Expression Omnibus under accession number GSE198621.

### Statistical Analysis

scRNA-seq was reproduced in three independent runs using DropSeq and 10x Chromium platforms. scRNAseq identified gene expression changes were validated in a separate cohort of identical UIR, UUO and control mice (n=4-6 per group). Effects of AHNAK knockdown on RPTEC EMT and mass migration were tested in 5 and 2 independent experiments, respectively (n=4 wells per group). P values were generated using one-way ANOVA with Bonferroni and Holm or Student’s t-test with *p<0.05 representing the statistically significant difference. Data are presented as individual values, mean ± SD.

## Results

### Ischemia/reperfusion and obstruction induced CKD models elicit dramatic proximal tubular loss, functional decline and novel cellular clusters

Two clinically relevant models – namely unilateral ischemia/reperfusion (UIR) and ureter obstruction (UUO)^38–40^ were established in C57Bl/6 male mice. Naive mice of the same strain, age and sex were used as controls. At day 28 post-injury, both models exhibited dramatic parenchymal reduction^41^ (Figure 1A). 10x Chromium scRNA-seq^31^ was carried out on the control, UIR and UUO left kidney suspensions. Single cell transcriptomes were analyzed, and kidney cell populations identified using unsupervised Uniform Manifold Approximation and Projection (UMAP) dimension reduction^32^. Control samples exhibited epithelial, glomerular and stromal clusters, including large proximal tubular population (*Slc34a1*), loop of Henle (*Slc12a1, Umod*), distal tubule (*Slc12a3*), collecting duct principal (*Aqp2*) and intercalated (*Atp6v1g3/0d2/1c2*) cells, endothelial (*Pecam1*) and podocyte (*Nphs1/2*) populations (Figure 1B, Supplemental figure 1A, Supplemental table 1). Controls also elicited small macrophage (*Lyz2, Ccl8*), T cell (*Cd3d/g*), B cell (*Cd19, Igkc*) and “Cell cycle” cluster expressing cell division (*Mki67, Ccnd1, Cdk1*) and DNA biosynthesis (*Topbp1, Top2a, Tyms*) related genes along with immune cell markers (*Ccr2, Wdfy4, Ifi205, Irf8*), which indicates that this population might represent proliferating immune cells (Supplemental figure 1B, Supplemental table 1).

**Figure 1.**
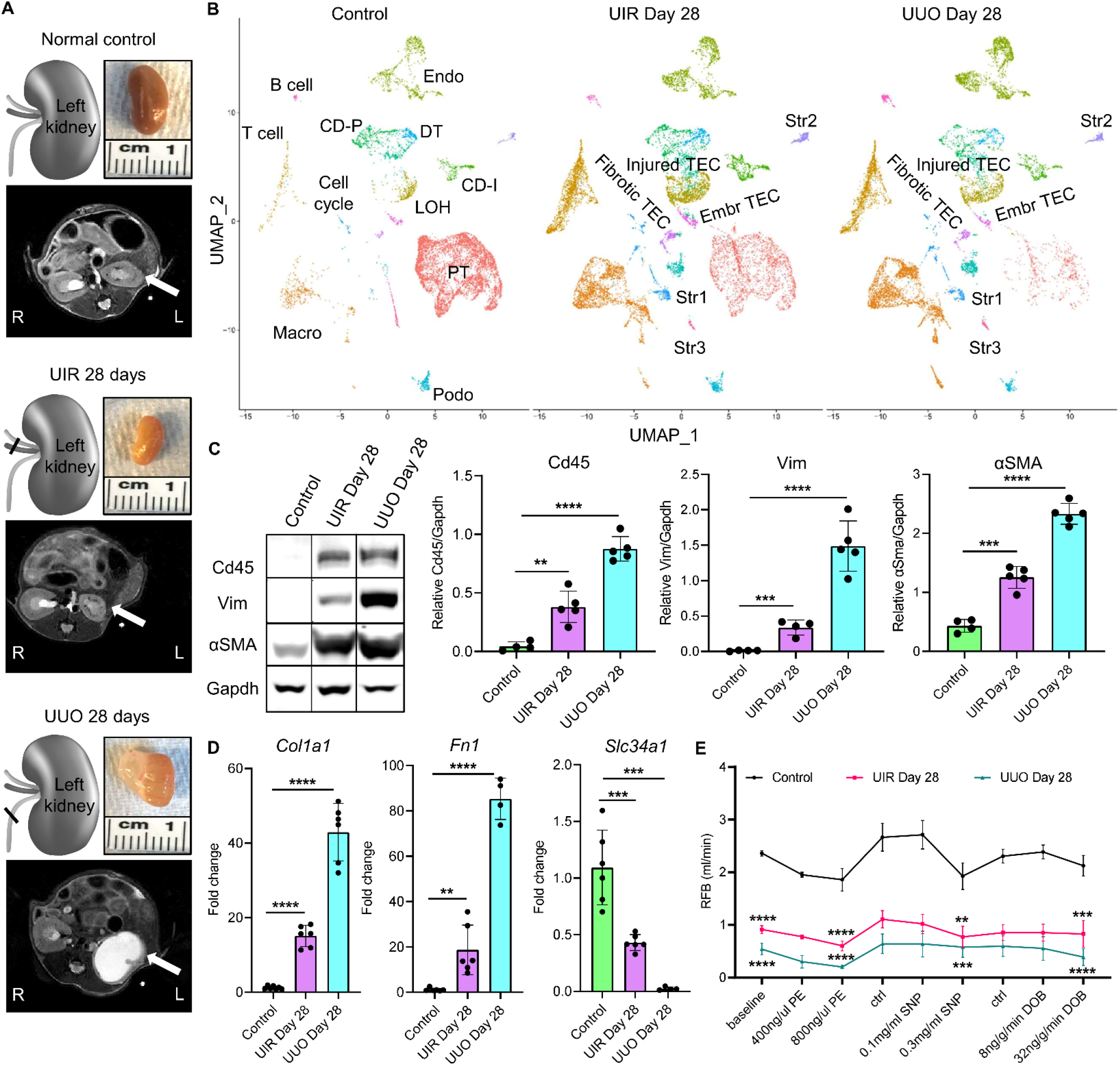
Ischemia/reperfusion and obstruction induced models of CKD elicit dramatic proximal tubular loss, kidney functional decline and novel cellular clusters. **(A)** Schemes of injury models, macroscopic and MRI images of normal control, UIR and UUO day 28 kidneys. Kidneys are pointed to with white arrows. R, right, L, left. **(B)** UMAPs show renal cell populations in the control, UIR and UUO kidneys (n=3-5 per group). Clusters are distinguished by different colors. PT, proximal tubules, LOH, loop of Henle, DT, distal tubule, CD-P, collecting duct principal, CD-I, collecting duct intercalated, Podo, podocytes, Endo, endothelial, Macro, macrophages, Str, stromal, TEC, tubular epithelial cells. **(C)** Representative images and quantifications of Western blots for fibrosis and inflammation markers, n=4-5 per group. **(D)** qPCR of fibrosis and proximal tubule markers, n=4-7 per group. **(E)** RBF at baseline, with vasoconstrictive (PE, phenylephrine), vasodilative (SNP, sodium nitroprusside) and inotropic agent (DOB, dobutamine). Agents administered at 0.1ul/min/gBW, Ctrl – control interval between treatments, n=3-4 per group, mean ± SD. **pValue ≤ 0.01, ***pValue ≤ 0.001, ****pValue ≤ 0.0001 compared to control, Student’s *t* test **(C)**, **(D)** and **(E)**.

scRNA-seq identified significant cellular landscape changes in UIR and UUO, including dramatic proximal tubule loss (Figure 1B). Both models elicited “Injured TECs” located between loop of Henle, distal tubules and principal collecting duct sub-cluster on the UMAP. Gene ontology (GO) analysis^42^ revealed apoptosis (*Anxa1, Xiap, Bcl10*), cytokine response (*Ifitm3, Ifi27, Cd47*), neutrophil mediated immunity (*S100a11, Lcn2, Ctss*), cell motility and actin filament organization (*Tmp1/4, Myo1d/e, Sptan1*), supramolecular fiber organization (*Vim, Flna, Krt8/19*) and epithelium development (*Cd24a, Sox4, Hoxb2/4*) related gene upregulation in this cluster (Supplemental figure 2A, Supplemental table 2). We also observed long-term injury induced macrophage, T-cell, B-cell and “Cell cycle” expansion (Figure 1B, Supplemental figures 3-5, Supplemental table 3). Moreover, scRNA-seq demonstrated remarkable interstitial expansion in both models, indicated by three distinctive stromal clusters. These changes were validated in independent control and fibrotic cohorts on RNA and protein levels (Figure 1, C and D). Moreover, both models elicited remarkable renal blood flow (RBF) decline with no systemic changes in heart rate or blood pressure (Figure 1E, Supplemental figure 6, A-C)^43^. Thus, we established two models exhibiting key CKD landmarks, including substantial renal functional decline, parenchymal loss, tubular injury, inflammation and fibrosis.

### scRNA-seq dissects the transcriptional identities and cellular origins of novel clusters contributing to kidney fibrotic remodeling

UIR and UUO resulted in mature *Slc34a1*-expressing proximal tubule loss and dramatic mesenchymal marker *Vim* elevation in the tubular, immune, and endothelial populations (Figure 2A). Single molecule fluorescent *in situ* hybridization (smFISH, with RNAscope) and Picrosirius Red staining validated remarkable *Slc34a1* decline along with peritubular and periglomerular expansion of *Vim*-expressing cells and ECM deposition in UIR and UUO cortex and medulla (Figure 2B). Thus, we used scRNA-seq to dissect the molecular identities of renal cell populations contributing to ECM production and fibrosis. GO analysis revealed ECM (*Mmp2/11/14, Fn1, Col1a1/2*), collagen fibril organization (*Lox, Loxl1/2/3, P3h1/h4*), wounding response (*Ext1, Fgf2, Flna*) and ossification (*Runx1, Bmp1/4, Bmpr1a/2*) related gene enrichment in the “Stromal 1” population (Figure 2C, Supplemental tables 1, 3). “Stromal 2” cells exhibited a strong muscle contraction signature, represented by actin filament process (*Acta2, Actb, Xirp1*), muscle structure development (*Mef2a/c/d, Myom1, Mylk, Myl6, Myh11, Smtn*) and contraction (*Tbx2, Pln, Utrn, Tpm1/2*) associated genes. In contrast, “Stromal 3” transcriptional signatures were enriched for cell migration, motility and locomotion (*Aamp, Mylk, Myh9, Amotl1/2*) along with vasculature development (*Vegfd, Angpt2, Pdgfa, Pdgfra/b*). We noted that while Stromal 1 and 2 populations expanded in the fibrotic kidneys, Stromal 3 cluster was equally present in the normal and injured conditions. Importantly, this observation was reproduced with an alternative Drop-seq platform^44^ (Supplemental figure 7, A-C). Importantly, doublet removal did not affect the control and fibrotic cellular landscapes, including three stromal clusters (Supplemental figure 8A). Overall, our analysis identified functional heterogeneity of quiescent and activated renal fibroblasts, evidenced by three distinctive populations with “secretory”, “contractile” and “migratory” signatures.

**Figure 2.**
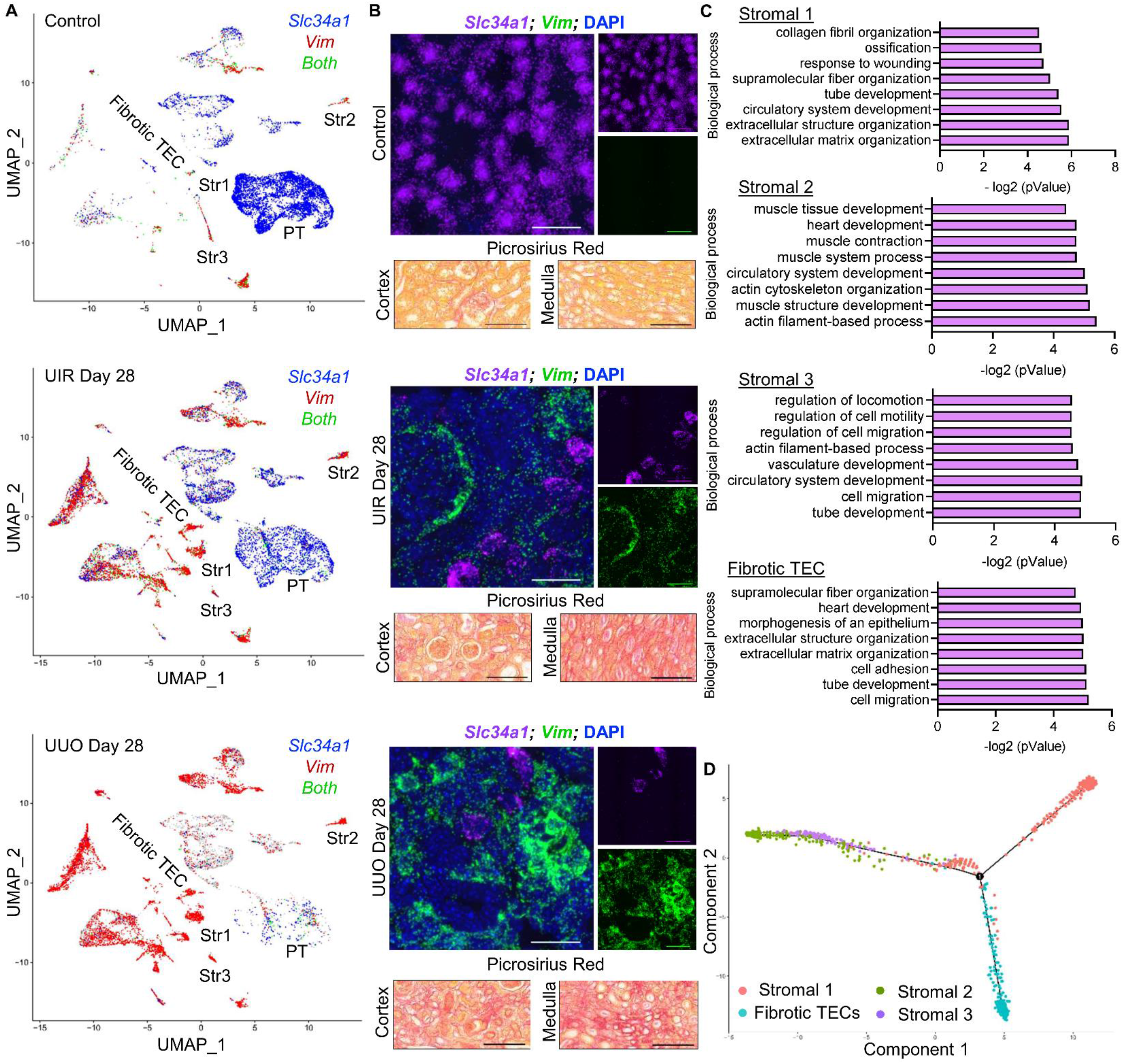
scRNA-seq dissects the transcriptional identities and cellular origins of novel clusters contributing to kidney fibrotic remodeling. **(A)** Feature plots revealing three distinctive stromal and fibrotic TEC clusters along with *Slc34a1* and *Vim* expression changes in control and fibrotic kidneys. **(B)** Upper panels: representative images of *Slc34a1* (purple), *Vim* (green) and DAPI (blue) RNAscope. Original magnification, ×60, 0.07 μm/px Nyquist zoom, maximal intensity projection from approximately 6-μm Z-stacks. Lower panels: representative images of Picrosirius Red staining showing increased ECM deposition in cortex and medulla (red color) of fibrotic kidneys compared to control. Original magnification, ×40, 0.16 μm/px zoom. **(C)** GO Biological process of “Stromal” and “Fibrotic TEC” marker genes vs other populations in control, UIR and UUO, – log2 (pValue). **(D)** Integrated trajectory analysis of control, UIR and UUO stromal and fibrotic TEC populations. Number 1 represents significant branch point of differentiation.

In addition to three stromal populations, both models exhibited novel “Fibrotic TECs” expressing both ECM and supramolecular fiber organization associated genes along with *Cdh16* encoding kidney specific epithelial marker cadherin 16^45^ (Figure 2C, Supplemental tables 1, 3). In addition to the mesenchymal transcriptional signature, fibrotic TECs elevated nephrogenic genes including *Cdh6*^46,47^ and *Osr2* (Supplemental figure 9, A and B). *Osr2* expression was correlated with specific epithelial-to-mesenchymal interaction sites in the developing kidney^36,48^. Thus, fibrotic TECs might represent a unique epithelial population that acquired a dedifferentiated mesenchymal phenotype in response to injury. However, integrated trajectory analysis found no evidence of significant EMT or endothelial-to-mesenchymal transition, consistent with the previous observations^18^ (Supplemental figure 10, A and B). Only a minor degree of transition was predicted between the stromal clusters as well as fibrotic TECs and “Stromal 1” population (Figure 2D). Integrated trajectory analysis also showed that stromal populations 2 and 3 clustered together, highlighting their transcriptional similarities (Supplemental figure 11A). Collectively, these results indicate that while injured TECs acquire remarkable dedifferentiated mesenchymal transcriptional signature, EMT does not significantly contribute to renal activated fibroblasts.

### Gucy1a3 specifically labels activated kidney fibroblasts in fibrotic CKD models

Next, we used scRNA-seq to identify markers which specifically label three activated kidney fibroblast clusters. Feature plots demonstrated that *Acta2* and *Col1a1* exhibited baseline and injury induced off-target expression in podocyte, immune, endothelial and tubular populations, including pro-fibrotic TECs (Figure 3A). Moreover, we found that *Acta2* was predominantly elevated in stromal clusters 2 and 3, while *Col1a1* only marked stromal 1 cells, illustrating transcriptional heterogeneity of activated renal fibroblasts. Additionally, we found that *Pdgfrb*, commonly used to label fibroblasts in multiple organs^49,50^, was not restricted to the stromal clusters (Supplemental figure 12A). Of note, we observed low-to moderate expression of *Gli1* (encoding GLI family zinc finger 1) in UIR and UUO stromal 1 cells (Supplemental figure 12B). Previous data showed that Gli1 labels perivascular mesenchymal stem cell (MSC)-like cells which differentiate into activated renal fibroblasts in response to the injury^51^. However, we demonstrated that *Gli1* does not thoroughly label all stromal clusters in the fibrotic kidneys.

**Figure 3.**
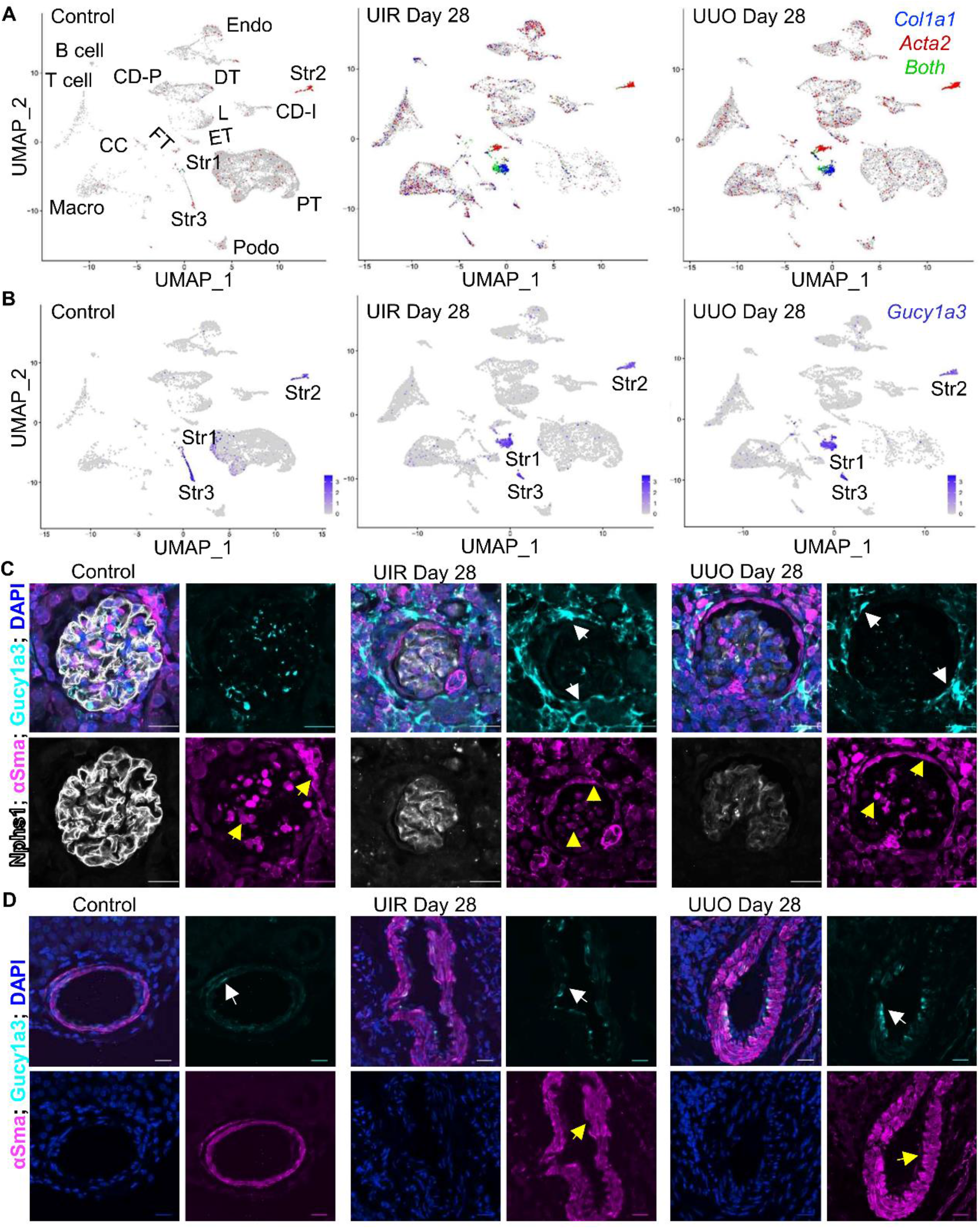
Gucy1a3 specifically labels activated kidney fibroblasts in fibrotic CKD models. **(A)** and **(B)** Feature plots demonstrating expression of historically used stromal markers *Acta2* and *Col1a1 vs* novel putative kidney activated fibroblast marker *Gucy1a3* in tubular, stromal, immune, endothelial and podocyte populations of normal and fibrotic kidneys. L, loop of Henle, FT, fibrotic TECs, ET, embryonic TECs, CC, cell cycle. **(C)** IF of αSma and Gucy1a3 podocyte and periglomerular expression in the normal and fibrotic kidneys. αSma expression in control, UIR and UUO Nphs1 -positive podocytes and inner periglomerular circle is shown with yellow arrows. Injury-induced outer periglomerular elevation of Gucy1a3 is shown with white arrows. **(D)** IF of αSma and Gucy1a3 expression in vascular SMCs of normal and fibrotic kidneys. Faint Gucy1a3 expression in control, UIR and UUO blood vessels is shown with white arrows. Remarkable injury induced αSma elevation in UIR and UUO blood vessels is highlighted with yellow arrows. Original magnification, ×60, 0.18 μm/px Nyquist zoom, maximal intensity projection from approximately 6-μm Z-stacks.

We found that *Gucy1a3* (a.k.a. *Gucy1a1*), encoding Guanylate Cyclase 1 Soluble Subunit Alpha 1, marked all three stromal clusters in both control and fibrotic kidneys (Figure 3B), with negligible off-target expression in endothelial, immune, podocyte and tubular compartments, including pro-fibrotic TECs, compared to *Vim, Acta2* and *Col1a1*. Immunofluorescent (IF) staining performed on an independent UIR, UUO and control cohort validated this finding. While Gucy1a3 was increased in periglomerular fibrotic areas of the UIR and UUO kidneys, it exhibited remarkably lower expression in Nphs1-positive glomerular podocytes compared to αSma (Figure 3C). Moreover, we showed that while injuries caused dramatic αSma elevation in the vascular SMCs, Gucy1a3 expression in the blood vessels of fibrotic kidneys remained at baseline (Figure 3D). This activated fibroblast specific expression was not changed after doublet removal (Supplemental figure 13A). Overall, our findings highlighted the potential of Gucy1a3 as a putative novel specific marker of activated kidney fibroblasts.

### Prolonged kidney injury elicits persistent nephrogenic signaling in the stromal and epithelial populations

scRNA-seq identified that both UIR and UUO not only caused mesenchymal changes in the injured epithelium, but also produced “Embryonic TECs”, co-expressing mature epithelial markers (*Cdh16, Slc34a1*) and renal developmental genes *Cdh6* and *Osr2* (Supplemental figure 9, A and B). This cluster was enriched for epithelium, kidney and urogenital system development (*Spry1, Cited2, Foxc1, Hnf1β, Pax2/8, Lgals3, Hoxa5/7, Hoxb2/4/5/7*) regulating genes^52^ (Figure 4A, Supplemental tables 1, 3). Furthermore, fibrotic kidneys thoroughly upregulated nephrogenic genes *Cd24a* and *Sox4*^48,53–55^ which we previously reported in AKI (Figure 4, B-D)^30^. scRNA-seq revealed remarkable elevation of both genes throughout the nephron tubule as well as in the fibrotic and embryonic TECs. *Cd24a* was also notably upregulated in UIR and UUO immune and cell cycle populations, reflecting its diverse immunological functions^56^. On the other hand, *Sox4* marked stromal clusters of both control and fibrotic kidneys, particularly Stromal 1 populations. We validated these findings using immunohistochemistry (IHC) and RNAscope, which demonstrated *Sox4* elevation in *Slc34a1* -and *Vim*-positive compartments (Figure 4, E and F). Collectively, we revealed that fibrotic kidneys exhibited enduring nephrogenic signaling in multiple populations.

**Figure 4.**
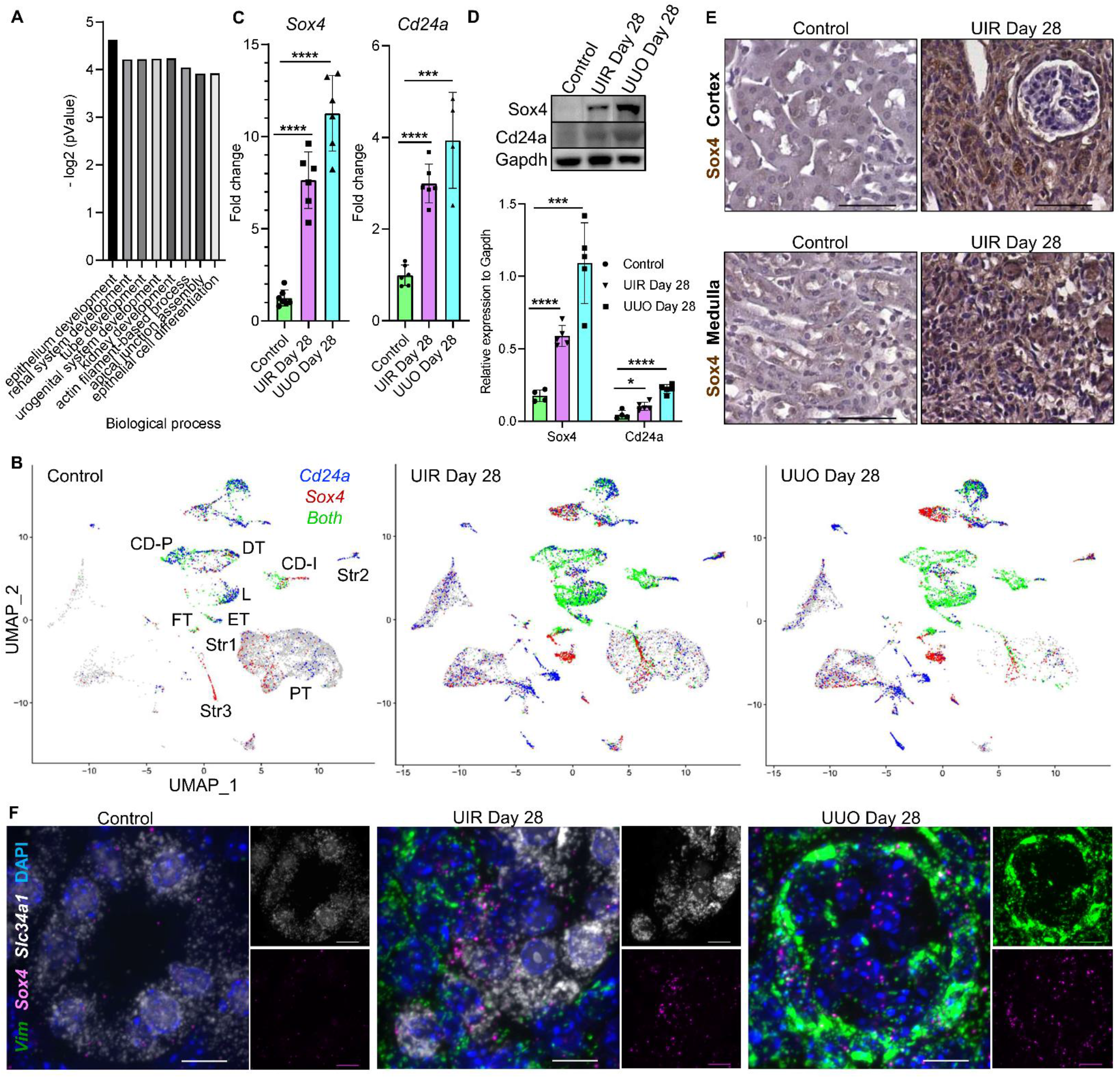
Prolonged kidney injury elicits persistent renal developmental signaling in the stromal and epithelial populations. **(A)** GO of “Embryonic TECs” marker genes vs other populations in control, UIR and UUO kidneys, - log2 (pValue). **(B)** Feature plots of *Sox4* and *Cd24a* expression in the tubular and stromal cells of control and fibrotic kidneys. **(C)** *Sox4* and *Cd24a* qPCR in control and fibrotic kidneys, n=4-7 per group. **(D)** Representative images and quantifications of Sox4 and Cd24a Western blots, n=4-5 per group. **(E)** Representative images of Sox4 IHC control and UIR kidneys. Original magnification, ×40, 0.16 μm/px zoom. **(F)** Representative RNAscope images demonstrating *Sox4* (pink) elevation in *Slc34a1* -positive (white) proximal tubular and *Vim*-positive (green) stromal compartments in UIR and UUO kidneys compared to mild expression in the control. DAPI (blue), original magnification, ×60, 0.05 μm/px Nyquist zoom, maximal intensity projection from approximately 6-μm Z-stacks. *pValue ≤ 0.05, ***pValue ≤ 0.001, ****pValue ≤ 0.0001 compared to control, Student’s *t* test, **(C)** and **(D)**.

### scRNA-seq dissects the novel cellular and molecular mechanisms of epithelial-to-stromal crosstalk in kidney fibrosis

Then, we asked whether fibrosis affects cell-to-cell communications in the kidney, particularly epithelial-to-stromal interactions. Ligand-receptor analysis predicted crosstalk between the control tubular segments via collagen, G protein, osteopontin, beta-2-microglobin, calmodulin, vascular endothelial growth factor, apolipoprotein and plasminogen signaling (Figure 5A, Supplemental table 4). Moreover, scRNA-seq predicted bidirectional interactions between resident stromal and epithelial populations via amyloid, calmodulin, calreticulin, Vegf, metalloproteinase inhibitor, phospholipase and transglutaminase pathways.

**Figure 5.**
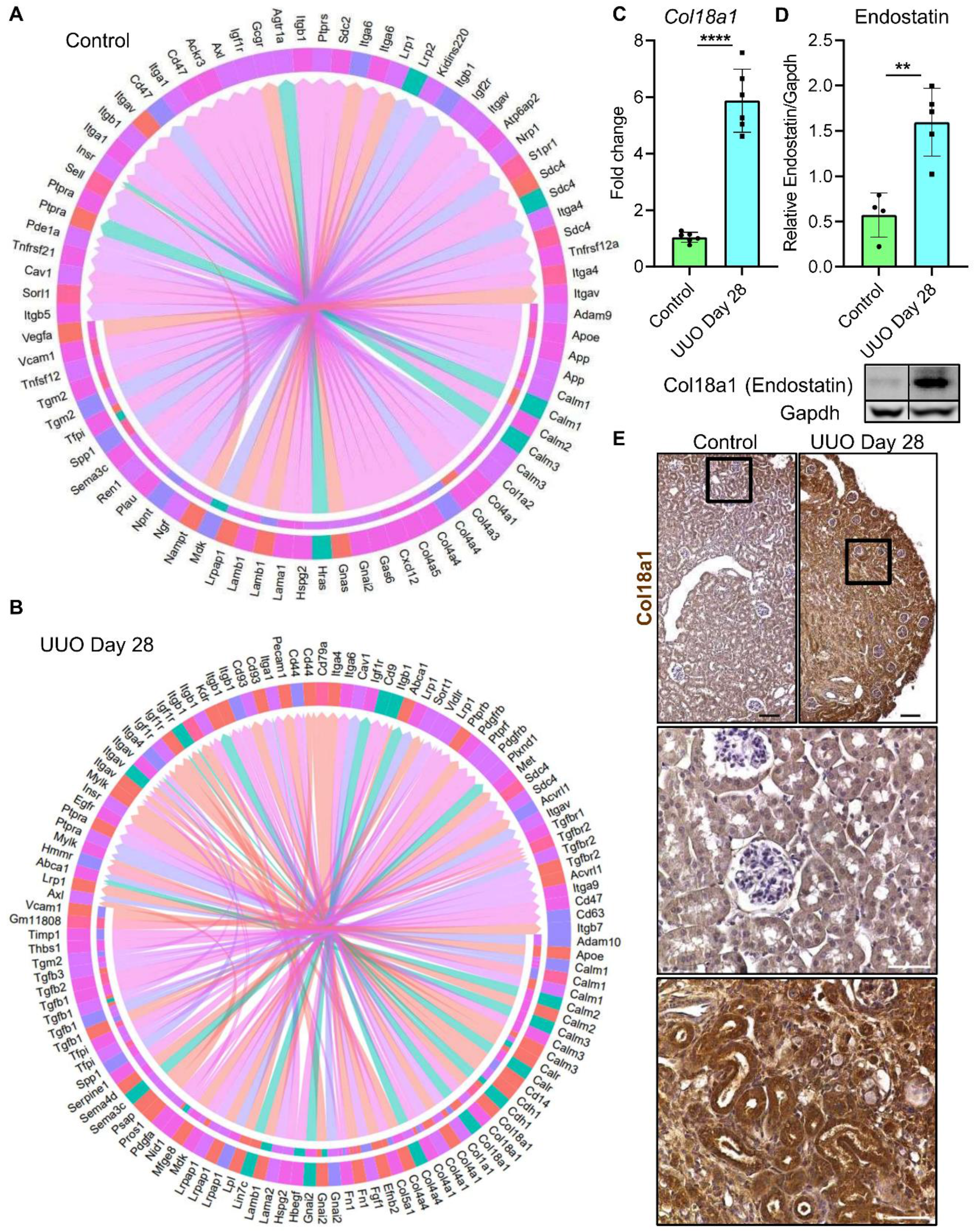
Long-term UUO induced kidney injury elicited pronounced epithelial-to-stromal crosstalk and elevation of Col18a1 signaling. **(A)** Circos plot of ligand-receptor interactions between proximal tubules (salmon), fibrotic TECs (heliotrope), embryonic TECs (magenta), stromal 1 (iris blue), stromal 2 (light slate blue) and stromal 3 (rose) clusters in control kidney. The populations producing putative ligand are shown at blunt end of the arrow; the populations producing putative receptor are highlighted with the color code line next to the ligand-producing cluster. Arrow points from the ligand-producing to receptor-producing populations. The names of all putative ligands and receptors with respect to the cell populations are available in Supplementary data 4. **(B)** Circos plot of ligand-receptor interactions between proximal tubules (salmon), fibrotic TECs (heliotrope), embryonic TECs (magenta), stromal 1 (iris blue), stromal 2 (light slate blue) and stromal 3 (rose) clusters in UUO kidney. **(C)** qPCR of *Col18a1* expression in control and UUO kidneys, n=6 per group. **(D)** Representative images and quantification of endostatin (*Col18a1* C-terminal product) Western blot, control and UUO kidneys, n=4-5 per group. **(E)** Representative images of Col18a1 IHC staining in control and UUO kidneys. Upper images - original magnification, ×10, 0.64 μm/px zoom; lower images – original magnification, ×40, 0.16 μm/px zoom into the upper images, area highlighted with black frames. **pValue ≤ 0.01, ****pValue ≤ 0.0001, compared to control, Student’s *t* test, **(C)** and **(D)**.

Both models exhibited dramatically increased cell-to-cell crosstalk, including bidirectional communications between UIR and UUO stromal and epithelial clusters (Figure 5B, Supplemental figure 14B). Specifically, “Stromal 1” and tubular clusters of both models interacted via *Spp1, Lcn2, Vim, Col4a1/3/4* and *Col18a1* signaling. Of note, activated Stromal 2 and 3, proximal tubules, loop of Henle and intercalated cells displayed upregulated *Tgfβ1* encoding Tgfβ ligand crucial for fibrosis. Corresponding Tgfβ receptor encoding genes (*Tgfβr1/2/3, Itgav, Itgb8*) were upregulated in all stromal and tubular clusters, including embryonic and fibrotic TECs, thus highlighting the bidirectional injury induced Tgfβ induction. Injured embryonic and fibrotic TECs also upregulated *Tgfβ2*, encoding another Tgfβ pathway ligand. scRNA-seq identified putative interactions between these clusters and other tubular segments along with immune, endothelial, stromal and podocyte populations via *Tgfβ2-Tgfβr1/2/3* and *Tgfβ2-Eng* ligand-receptor pairs. We also found that embryonic and fibrotic TECs might influence each other via many additional renal development and fibrosis regulating pathways including *Fn1-Itgb1, Bmp4-Bmpr1a/2, Jag1-Notch1/2/3* (Supplemental table 4).

Ligand-receptor analysis also demonstrated that fibrotic remodeling elicited elevated epithelial-to-stromal signaling via *Vim-Cd44* and *Col1a1/2-Itgb1/Itgav/Cd36/44* pathways, highlighting the active role kidney epithelium plays in pro-fibrotic phenotype induction. Moreover, scRNA-seq newly identified *Col18a1-Itgb1/a4/Gpc1/4* pathway in advanced kidney injury induced epithelial-to-stromal crosstalk (Figure 5B, Supplemental figure 14B, Supplemental table 4).We previously implicated *Col18a1*^57^ in AKI induced epithelial-to-stromal interactions^30^. *Col18a1* RNA and protein elevation was validated in both kidney fibrosis models (Figure 5 and Supplemental figure 14, C-E, Supplemental figure 15A). Overall, our findings predict bidirectional epithelial-to-stromal induction and highlight Col18a1 as a putative novel mediator of these interactions in kidney fibrosis.

### scRNA-seq reveals novel TEC gene expression signatures in kidney fibrosis

Since we showed strong dedifferentiated mesenchymal signature in the chronically injured epithelium, we asked which novel mediators might regulate these maladaptive changes. UIR and UUO proximal tubules and pro-fibrotic TECs elevated genes regulating actin filament binding (*Actn1, Myh9, Tpm1/3/4*), integrin binding (*Itgav, Itgb1, Myh9*), ECM and collagen binding (*Pdgfb, Lgals1/3*), ECM structure and binding (*Col1a1/2, Col4a1/2/3/4/5/6, Col6a1/2*), cadherin binding (*Ahnak, Myh9, Tagln2*), cell adhesion molecule binding (*Ahnak, Myh9, Spp1*) and structural molecule activity (*Vim, Tubb, Ahnak*) biological processes (Supplemental figure 16, A and B, Supplemental table 5). We identified and validated the persistent *Spp1* (encoding osteopontin) elevation in both kidney fibrosis models, which we previously reported in AKI^30^ (Supplemental figure 16C). Our data also revealed persistent elevation of many novel AKI-inducible genes we previously reported^30^, including *Myh9* and *Sh3bgrl3* (Supplemental figure 17, A-C). Remarkably, both CKD models exhibited enduring upregulation of *Ahnak*, encoding Neuroblast Differentiation-Associated Protein AHNAK (a.k.a. desmoyokin). Normal kidney moderately expressed *Ahnak*, while UIR and UUO resulted in remarkable *Ahnak* induction in the proximal tubules, fibrotic and embryonic TECs, and three activated fibroblast clusters (Figure 6A, Supplemental figure 16, A and B). Quantitative validation confirmed significant Ahnak increase in both fibrosis models (Figure 6, B and C, Supplemental figure 17A). Spatial analysis also verified remarkable Ahnak elevation in the stromal and tubular compartments of the fibrotic kidneys, particularly in E-cadherin positive TECs and Aqp1-positive proximal tubules (Figure 8, D-F).

**Figure 6.**
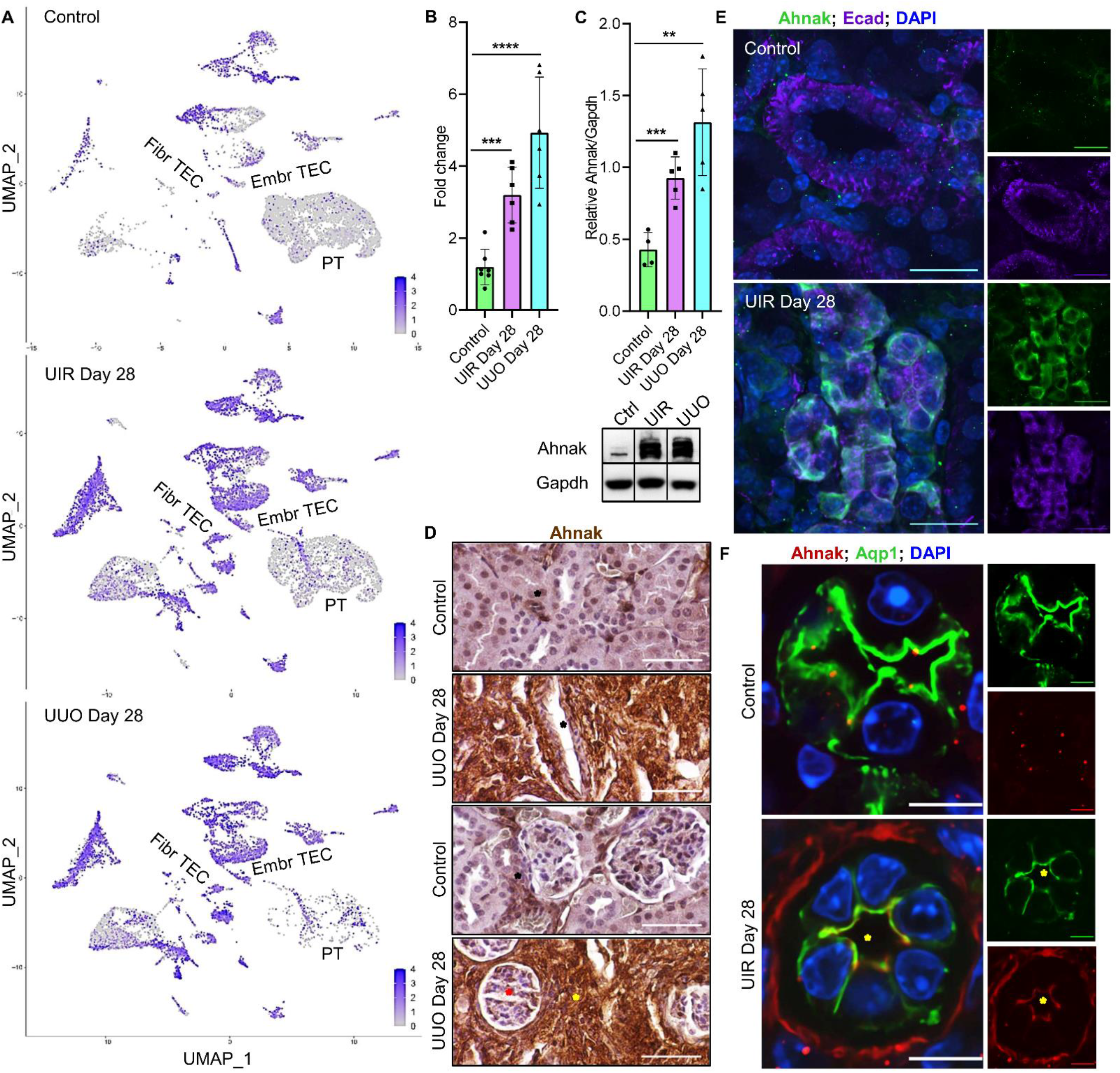
Ahnak elevation persists in tubular epithelial cells of fibrotic kidneys, including proximal tubules. **(A)** Feature plots of *Ahnak* in control and fibrotic kidneys. Gene expression levels are color-coded. **(B)** qPCR of *Ahnak* in control and fibrotic kidneys, n=4-7 per group. **(C)** Representative images and quantification of Ahnak Western blot, control, UIR and UUO kidneys, n=4-5 per group. **(D)** Representative images of Ahnak IHC in control and UUO kidneys. Mild stromal expression in the normal kidney is pointed with black asterix; vascular, podocyte and ubiquitous tubular elevation in UUO kidneys is highlighted with black, red and yellow asterix, respectively. Original magnification, ×40, 0.16 μm/px zoom. **(E)** Representative images of injury-induced Ahnak (green) elevation in Ecad-positive (purple) kidney tubules. IF staining, original magnification, ×60, 0.05 μm/px Nyquist zoom, maximal intensity projection from approximately 6-μm Z-stacks. **(F)** Representative images of injury-induced Ahnak (red) elevation on apical surfaces of Aqp1-positive (green) kidney proximal tubules (shown with yellow asterix). IF staining, original magnification, ×60, 0.05 μm/px Nyquist zoom, maximal intensity projection from approximately 6-μm Z-stacks. **pValue ≤ 0.01, ***pValue ≤ 0.001, ****pValue ≤ 0.0001 compared to control, Student’s *t* test, **(B)** and **(C)**.

### AHNAK knockdown exacerbates TGFβ response in primary human renal proximal tubular epithelial cells (RPTECs)

Our current and previous studies demonstrated remarkable AKI^30^ and CKD induced tubular epithelial elevation of Ahnak, including in proximal tubules and fibrotic TECs. Other groups showed that Ahnak regulates migration and invasion and potentiates TGFβ induced EMT in human keratinocytes and murine melanoma cells, but the role of Ahnak in kidney fibrosis is unknown^58^. Thus, we questioned whether Ahnak plays a mechanistic role in maladaptive renal epithelial pro-fibrotic changes and established a TGFβ dependent EMT assay in primary human RPTECs treated with AHNAK specific or non-targeting siRNA. IF revealed AHNAK expression in the cytoplasm and membrane of control, vehicle (Vh) and siScramble treated RPTECs and spindle cell shape changes in response to TGFβ (Figure 7A). siAHNAK+Vh and siAHNAK+TGFβ groups exhibited no detectable AHNAK expression. Western blot verified significant loss of AHNAK in both siAHNAK+Vh and siAHNAK+TGFβ RPTECs (Figure 7, B and C, Supplemental figure 18A). Then, we assessed EMT markers and found that AHNAK silencing caused substantial FN, αSMA and N-cadherin (NCAD) overexpression both with and without TGFβ in RPTECs (Figure 7, B and C). Prolonged AHNAK ablation resulted in significantly impaired proximal tubular (Gamma-Glutamyltransferase 1, GGT1) and pan-epithelial (ECAD) markers expression, as shown by both qPCR and Western blot (Figure 7, B-E). Moreover, IF revealed that AHNAK knockdown elicited disrupted membrane expression of epithelial marker Tight junction protein 1 (a.k.a. ZO1) with and without TGFβ (Figure 7F). Overall, we found that AHNAK knockdown promoted baseline mesenchymal changes and exacerbated TGFβ response in the RPTEC *in vitro* model of EMT.

**Figure 7.**
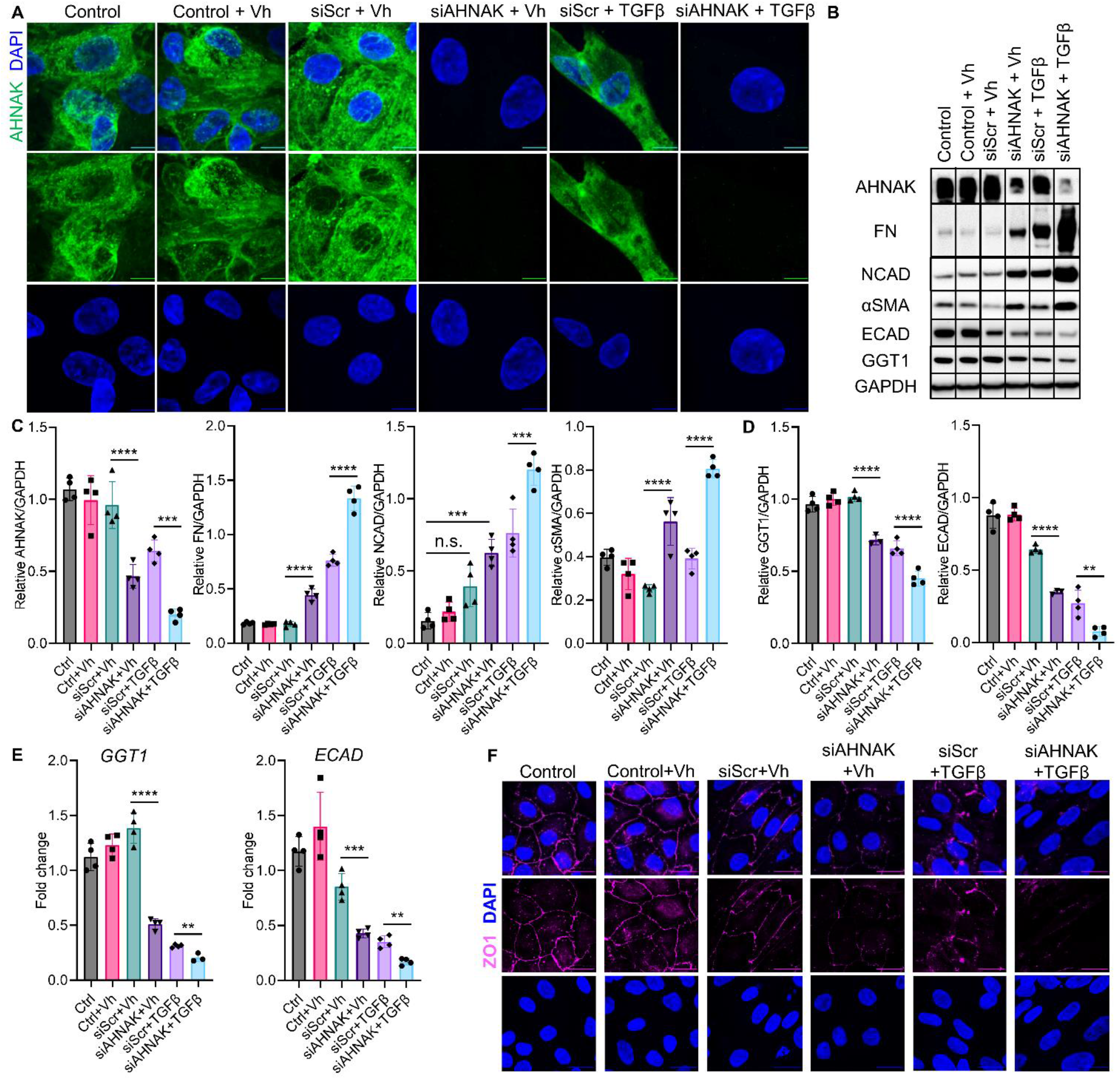
AHNAK knockdown exacerbates EMT in primary human RPTECs. **(A)** Representative images of AHNAK ICC in primary human RPTECs at the baseline and treated with Vh or TGFβ. 48 hours TGFβ treatment caused EMT outlined by spindle shape changes in RPTECs. siAHNAK administered groups exhibited no detectable AHNAK expression with and without TGFβ. AHNAK, green, DAPI, blue. Original magnification, ×60, 0.06 μm/px Nyquist zoom, maximal intensity projection from approximately 6-μm Z-stacks. **(B-D)** Representative images **(B)** and quantification of Western blot for AHNAK, EMT markers FN, NCAD and αSMA, n=4 per group **(C)**, epithelial markers ECAD and GGT1, n=3-4 per group **(D)** in RPTECs. AHNAK and EMT markers are assessed 48 hours post-TGFβ treatment, epithelial markers - following prolonged 72 hours TGFβ treatment. The effects of AHNAK knockdown were observed in 5 independent sets of experiments. **(E)** qPCR of *Ecad* and *Ggt1* in RPTECs 72 hours post-TGFβ, n=3-4 per group. **(F)** Representative images of epithelial marker ZO1 ICC in primary human RPTECs at the baseline and treated with Vh or TGFβ. siAHNAK treatment caused decreased membrane expression of ZO1 both with and without TGFβ. ZO1, pink, DAPI, blue. Original magnification, ×60, 0.07 μm/px Nyquist zoom, maximal intensity projection from approximately 6-μm Z-stacks. **pValue ≤ 0.01, ***pValue ≤ 0.001, ****pValue ≤ 0.0001, n.s., non-significant, comparisons shown with black lines, one-way ANOVA **(C, D)**, Student’s *t* test **(E)**.

### AHNAK affects migration and alters p38, p42/44, pAKT, BMP and MMP signaling in the primary human RPTEC in vitro model of EMT

Since AHNAK mediated EMT related changes in RPTECs, we asked whether it affects their functional capacities. Analysis of multiple time points over a 24-hour period demonstrated impaired migration in siAHNAK groups compared to siScramble RPTECs (Figure 8, A and B). Next, we evaluated protein levels of TGFβ canonical and non-canonical pathway members 1-hour post-induction and found that AHNAK deficiency caused decreased levels of phosphorylated p44/42 and AKT (Figure 8C). On the contrary, early p38 phosphorylation was increased in AHNAK depleted RPTECs. AHNAK ablation did not affect SMAD2/3 phosphorylation which suggests that it might act through the non-canonical TGFβ pathway. Remarkably, even early after TGFβ administration, AHNAK knockdown caused significant decrease of bone morphogenetic protein-7 (BMP7), known for anti-fibrotic effects in CKD^59^. Then, we examined later stages of TGFβ response and found that AHNAK ablation elicited increased p44/42 phosphorylation in both Vh- and TGFβ-treated conditions (Figure 8D). Moreover, AHNAK deficient RPTECs exhibited persistent BMP7 decline in both siAHNAK+Vh and siAHNAK+TGFβ groups. AHNAK silencing also reduced levels of *MMP2* encoding matrix metalloprotease-2 involved in collagen degradation (Figure 8E)^60^. Collectively, our data identified that AHNAK modulates EMT in primary human RPTECs via non-canonical TGFβ pathway, BMP and MMP signaling.

**Figure 8.**
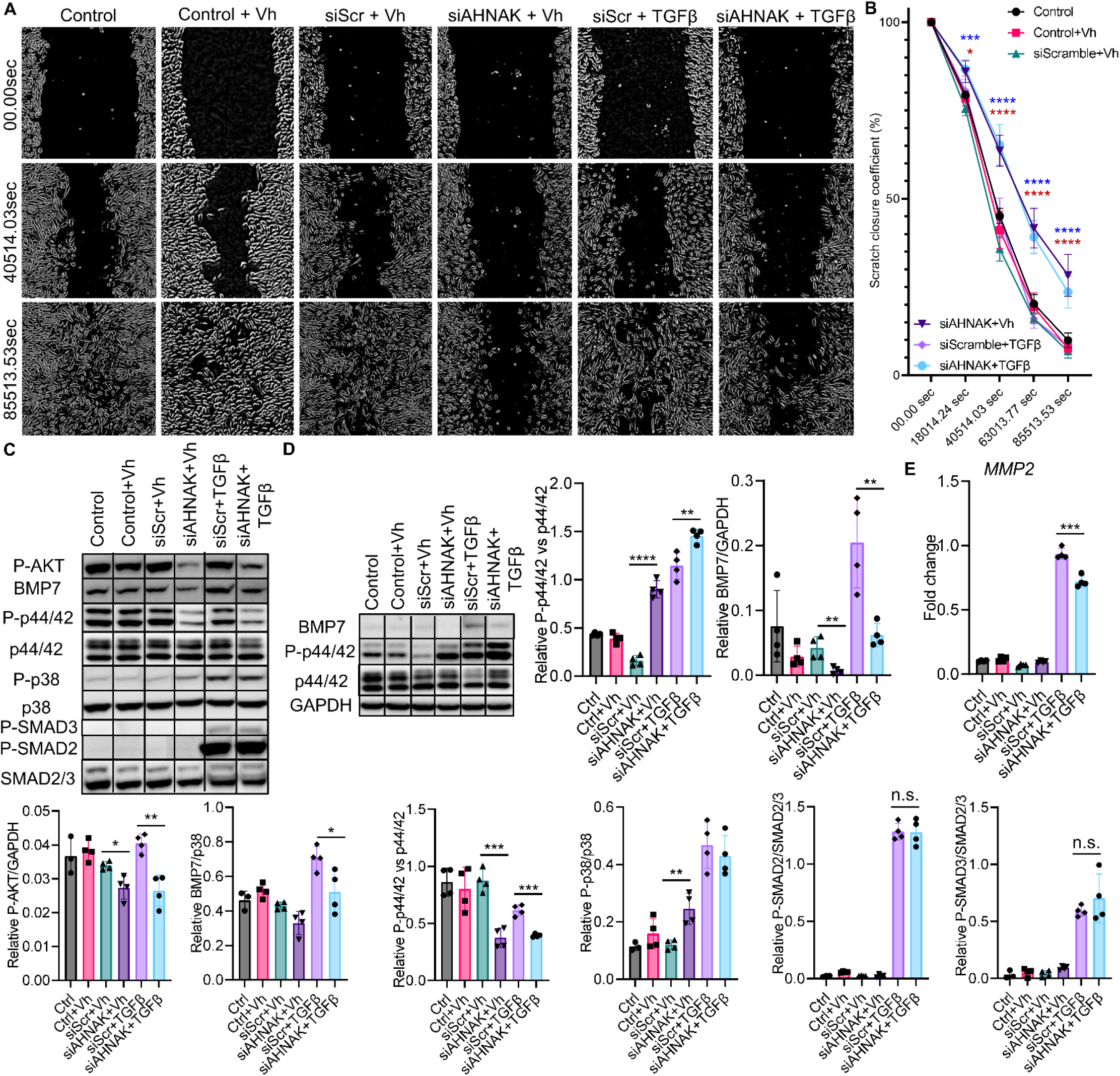
AHNAK affects migration and alters p38, p42/44, pAKT, BMP and MMP signaling in the primary human RPTEC *in vitro* model of EMT. **(A)** Representative images of 24 hour mass migration assay (00.00sec – first time point, 40514.03sec – second time point post-scratch, 85513.53sec – last time point post-scratch) in primary human RPTECs treated with siScramble or siAHNAK with or without TGFβ. Original magnification, ×40, 0.65 μm/px zoom. This observation was reproduced in two independent experiments. **(B)** Quantification of the mass migration assay from (A). 00.00 sec represents beginning of the experiment. Migration is assessed as scratch closure coefficient (%) at 18014.24, 40514.03, 63013.77, 85513.53 sec post-scratch and TGFβ administration. N=4 per group, pValues are color-coded: siScramble+Vh vs siAHNAK+Vh in blue, siScramble+TGFβ vs siAHNAK+TGFβ as red. **(C)** Representative images and quantification of Western blots for P-p38, P-p44/42, P-AKT, P- SMAD2/3, BMP7 at 1 hour post-TGFβ administration, n=3-4 per group. **(D)** Representative images and quantification of Western blots for P-p44/42 and BMP7 assessed at 48 hours post-TGFβ administration, n=4 per group. **(E)** RT-qPCR of *MMP2* in RPTECs 48 hours post-TGFβ, n=4 per group. N.s., non-significant, *pValue ≤ 0.05, **pValue ≤ 0.01, ***pValue ≤ 0.001, ****pValue ≤ 0.0001, comparisons shown with black lines, one-way ANOVA **(B)**, Student’s *t* test **(C-E)**.

## Discussion

We implemented a powerful, unbiased scRNA-seq approach to examine the molecular and cellular landscapes of advanced kidney injury. We combined bioinformatics analysis with comprehensive validation and mechanistic studies to dissect two crucial contributors to maladaptive kidney injury response and fibrotic remodeling – aberrant fibroblast activation and enduring pro-fibrotic phenotype in TECs.

Despite recent advances in understanding the molecular and cellular landscapes of normal and diseased human and murine kidney^30,49,61–65^, renal quiescent and activated fibroblasts remain an elusive population. Kuppe et al^49^ used spatial transcriptomics and genetic fate tracing to examine human CKD and murine UUO induced renal fibrosis. However, defining renal activated fibroblasts via ECM genes or *Pdgfrβ* might not allow for distinction between the tubular and stromal cells contributing to renal fibrosis. Our scRNA-seq utilizing two independent platforms reproducibly identified three novel fibroblast clusters exhibiting “secretory”, “contractile” and “migratory” transcriptional signatures in the normal and fibrotic kidneys. We discovered that Gucy1a3^66^ was specifically upregulated in three adult stromal clusters with minimal off-target expression. Future research will further test Gucy1a3 potential for specific activated renal fibroblast targeting in rodent and human CKD.

Previous studies^30,63,67–70^ revealed that failed TEC repair elicits pro-fibrotic phenotypes promoting AKI- to-CKD progression. Our analysis predicted increased intercellular communications in kidney fibrosis and corroborated the active role of TECs in the fibroblast activation. Furthermore, our studies identified Ahnak, previously reported in AKI^30^, as a mechanistic regulator of maladaptive pro-fibrotic changes in TECs. Primary human *in vitro* model of EMT revealed that AHNAK knockdown promoted mesenchymal changes and attenuated epithelial phenotype in RPTECs with and without TGFβ. Moreover, we identified the downstream signaling pathways underlying exacerbated TGFβ response in AHNAK depleted RPTECs, including anti-fibrotic BMP7 signaling.

Overall, our novel findings might hold a remarkable potential for the field, offering a thorough single cell resolution atlas of molecular and cellular changes featuring kidney fibrosis. Identification of Gucy1a3 as a novel specific kidney fibroblast marker, and Ahnak as an injury target, might improve our approach to understanding and halting kidney fibrosis respectively.

## Author contributions

VRM, MA, KS, AP, DV, IVA, JMK, QM, SP and PD designed experiments, analyzed results, and reviewed the manuscript. VRM, AP, KS, SMC, IVA, DML and MLN conducted experiments and acquired data. VRM, MA and PD performed final manuscript review.

## Acknowledgements

This work was supported by grants RO1 HL13395 and P50 DK096418 to PD.

## Data sharing

scRNA-seq data were deposited at the Gene Expression Omnibus under accession number GSE198621.

## Supplemental material table of contents

### Supplemental tables

**Supplemental table 1:** Marker genes used to identify cell types in the scRNA-seq data.

**Supplemental table 2:** Genes elevated or downregulated in UIR and UUO “Injured TECs” compared to control cluster.

**Supplemental table 3:** Cell type markers corresponding to Control, UIR and UUO individual heatmaps.

**Supplemental table 4:** Ligand-receptor interactions analysis in control, UIR and UUO kidneys.

**Supplemental table 5:** Genes elevated or downregulated in UIR and UUO “Proximal tubules”, “Embryonic TECs” and “Fibrotic TECs” compared to corresponding control clusters.

### Supplemental figures

**Supplemental figure 1.** scRNA-seq identifies kidney cellular clusters of control, UIR and UUO mice.

**Supplemental figure 2.** GO biological process analysis identifies transcriptional signature of “Injured tubular” clusters of fibrotic kidneys.

**Supplemental figure 3.** Heatmap reveals cellular landscapes and marker genes of control kidney.

**Supplemental figure 4.** Heatmap reveals cellular landscapes and marker genes of UIR Day 28 kidney.

**Supplemental figure 5.** Heatmap reveals cellular landscapes and marker genes of UUO Day 28 kidney.

**Supplemental figure 6.** Invasive hemodynamic reveals no significant systemic cardiovascular function changes caused by UIR and UUO compared to control.

**Supplemental Figure 7.** Drop-seq reveals molecular and cellular landscapes of kidney fibrosis in an independent cohort of UIR and UUO mice.

**Supplemental Figure 8.** Doublet removal does not affect scRNA-seq identified cellular landscapes in the normal and fibrotic kidneys.

**Supplemental Figure 9.** scRNA-seq reveals transcriptional identity of fibrotic TECs.

**Supplemental figure 10.** Integrated trajectory analysis shows little evidence of EMT or endothelial-to-mesenchymal transition in advanced renal injury.

**Supplemental figure 11.** scRNA-seq integrated trajectory analysis reveals transcriptional identities of kidney ECM producing stromal and tubular clusters.

**Supplemental figure 12.** scRNA-seq demonstrates stromal markers *Pdgfrb* and *Gli1* expression in normal and fibrotic kidneys.

**Supplemental figure 13.** Gucy1a3 specifically labels activated kidney fibroblasts in the control and fibrotic kidney scRNA-seq datasets after doublet removal.

**Supplemental figure 14.** scRNA-seq dissects the novel cellular and molecular mechanisms of crosstalk between the stromal and epithelial cells in the UIR model of kidney fibrosis.

**Supplemental figure 15.** scRNA-seq demonstrates *Col18a1* expression in normal and fibrotic kidneys.

**Supplemental figure 16.** scRNA-seq reveals novel gene expression signatures in tubular epithelial cells in kidney fibrosis.

**Supplemental figure 17.** CKD models elicit enduring elevation of AKI-inducible genes *Myh9* and *Sh3bgrl3*.

**Supplemental figure 18.** Western blot validates UIR and UUO inducible Ahnak elevation in fibrotic kidneys.

**Supplemental figure 19.** Western blot validates AHNAK siRNA knockdown.

