## Supplemental data for "Single-cell transcriptomic profiling of kidney fibrosis identifies a novel specific fibroblast marker and putative disease target"

**Supplemental figure 18.** Western blot validates UIR and UUO inducible Ahnak elevation in fibrotic kidneys.

**Supplemental figure 19.** Western blot validates AHNAK siRNA knockdown.

**Supplemental figure 1. scRNA-seq identifies kidney cellular clusters of control, UIR and UUO mice. (A)** Combined control, UIR and UUO feature plots demonstrate markers used to identify renal cell populations. Expression levels are color-coded. **(B)** GO Biological process of “Cell cycle” marker genes vs other populations in control, UIR and UUO, – log2 (pValue).

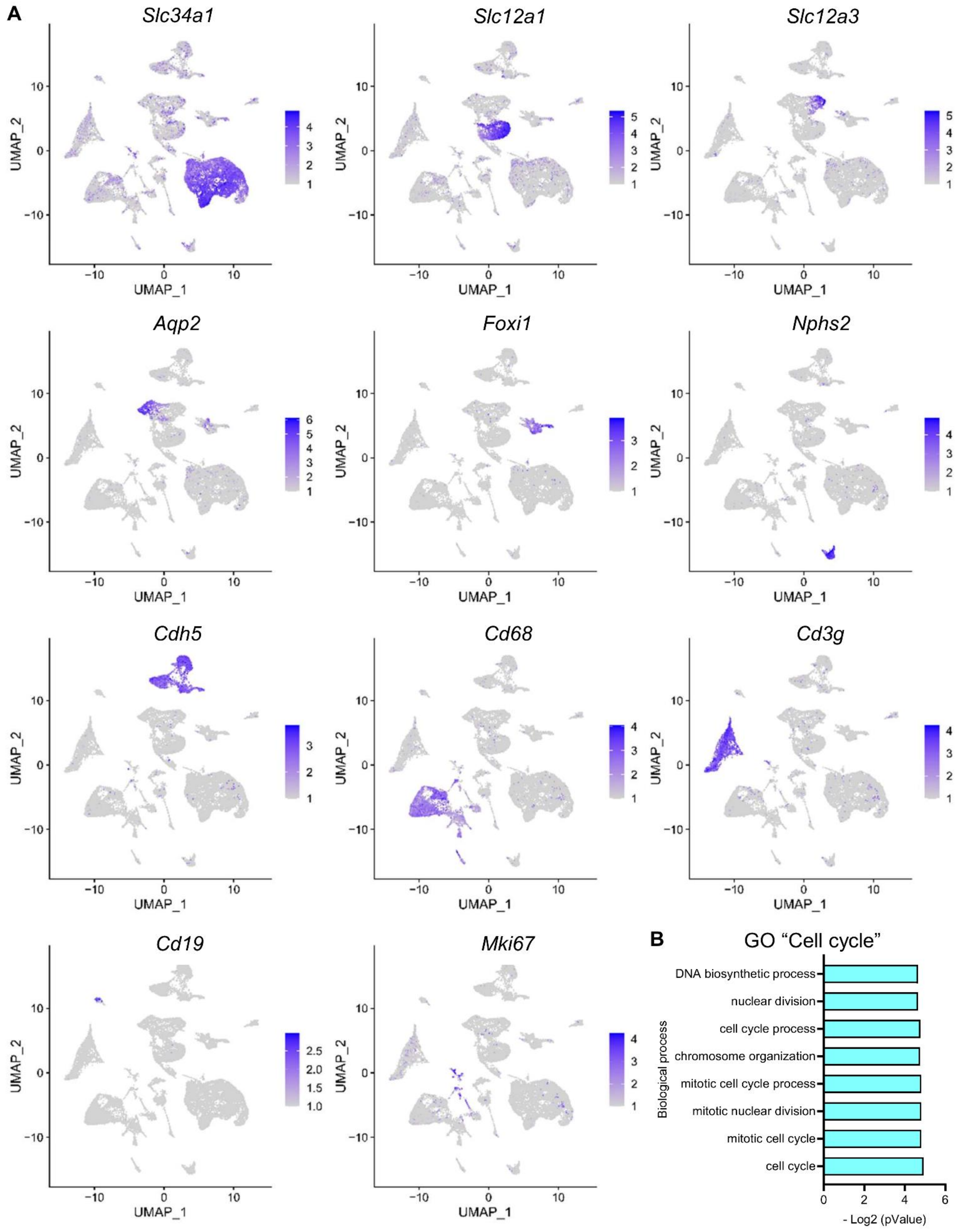

**Supplemental figure 2. GO biological process analysis identifies transcriptional signature of "Injured tubular" clusters of fibrotic kidneys. (A)** GO biological process of genes elevated in UIR and UUO "Injured tubular" clusters vs control, – log2 (pValue).

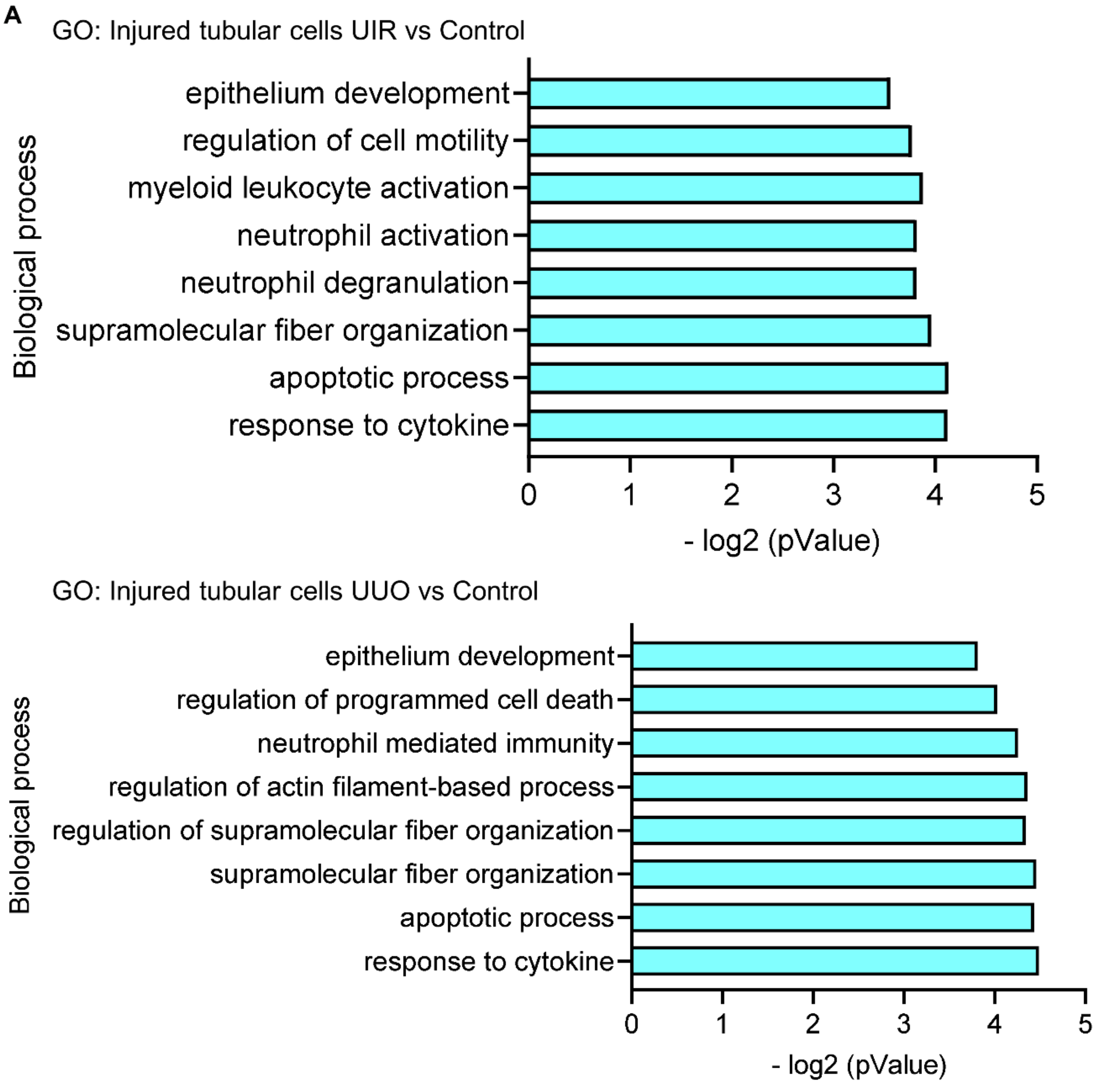

**Supplemental figure 3. Heatmap reveals cellular landscapes and marker genes of control kidney. (A)** Heatmap shows the relative marker gene expression in control renal cell types. Yellow color represents expression level above the mean, black color represents the mean, and purple/blue represents expression level below the mean. The heatmap shows genes elevated in renal cell populations relative to each other, based on the z-score. Representative marker genes are listed next to the heatmap. Complete list of control marker genes is attached as Supplemental table 3.

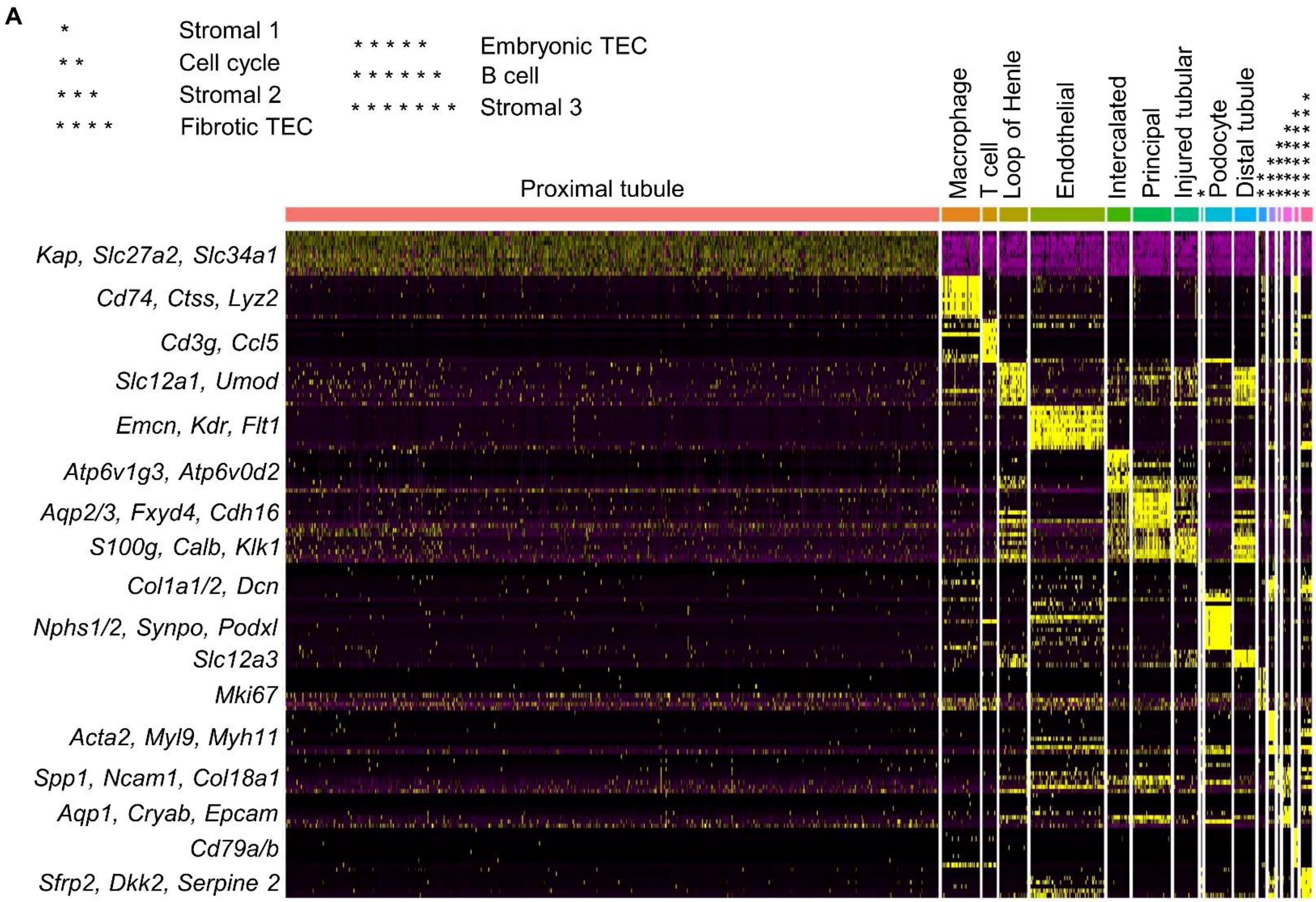

**Supplemental figure 4. Heatmap reveals cellular landscapes and marker genes of UIR Day 28 kidney. (A)** Heatmap shows the relative marker gene expression in UIR Day 28 renal cell types. Yellow color represents expression level above the mean, black color represents the mean, and purple/blue represents expression level below the mean. The heatmap shows genes elevated in renal cell populations relative to each other, based on the z-score. Representative marker genes are listed next to the heatmap. Complete list of UIR Day 28 marker genes is attached as Supplemental table 3.

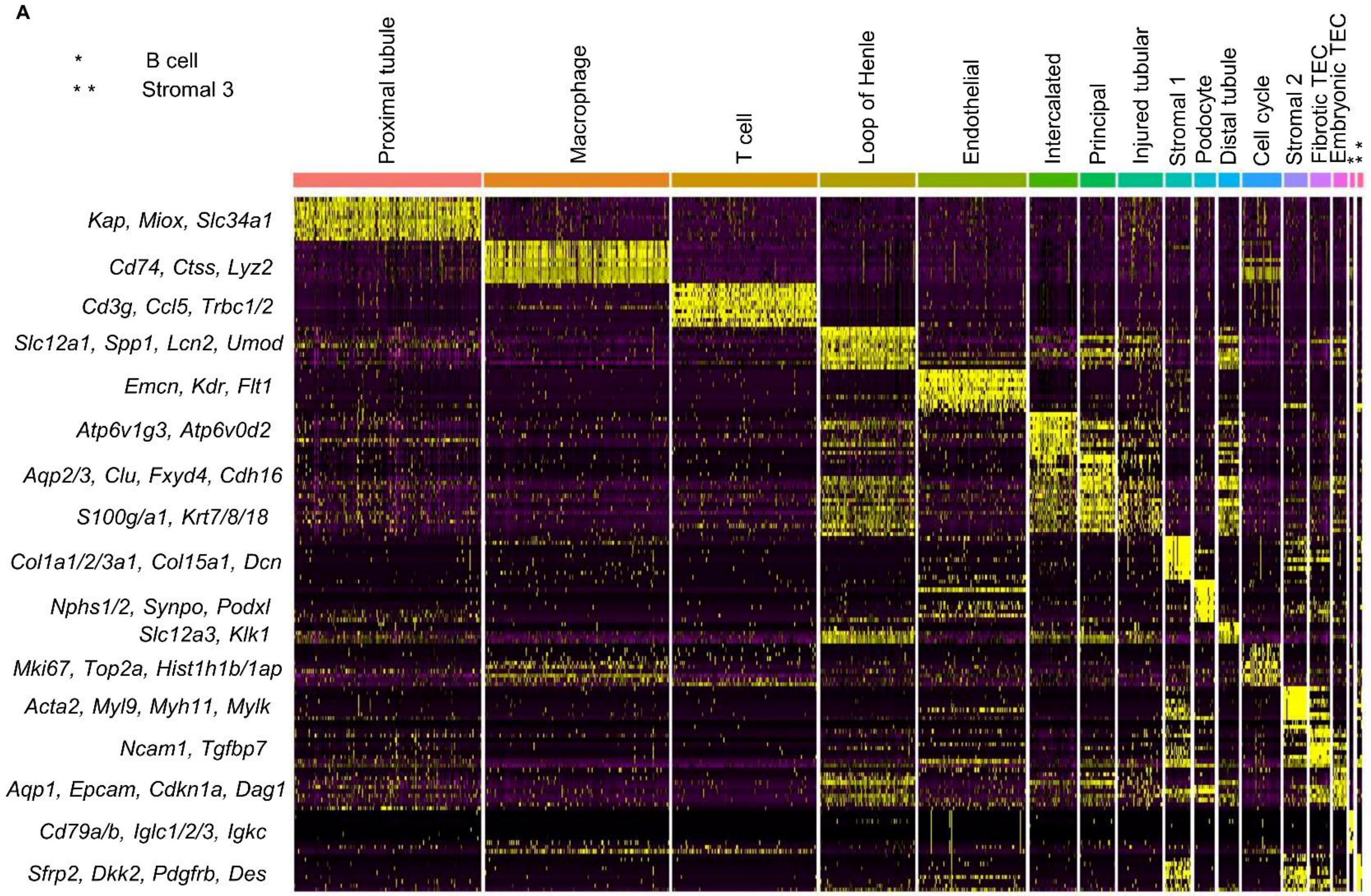

**Supplemental figure 5. Heatmap reveals cellular landscapes and marker genes of UUO Day 28 kidney. (A)** Heatmap shows the relative marker gene expression in UUO Day 28 renal cell types. Yellow color represents expression level above the mean, black color represents the mean, and purple/blue represents expression level below the mean. The heatmap shows genes elevated in renal cell populations relative to each other, based on the z-score. Representative marker genes are listed next to the heatmap. Complete list of UUO Day 28 marker genes is attached as Supplemental table 3.

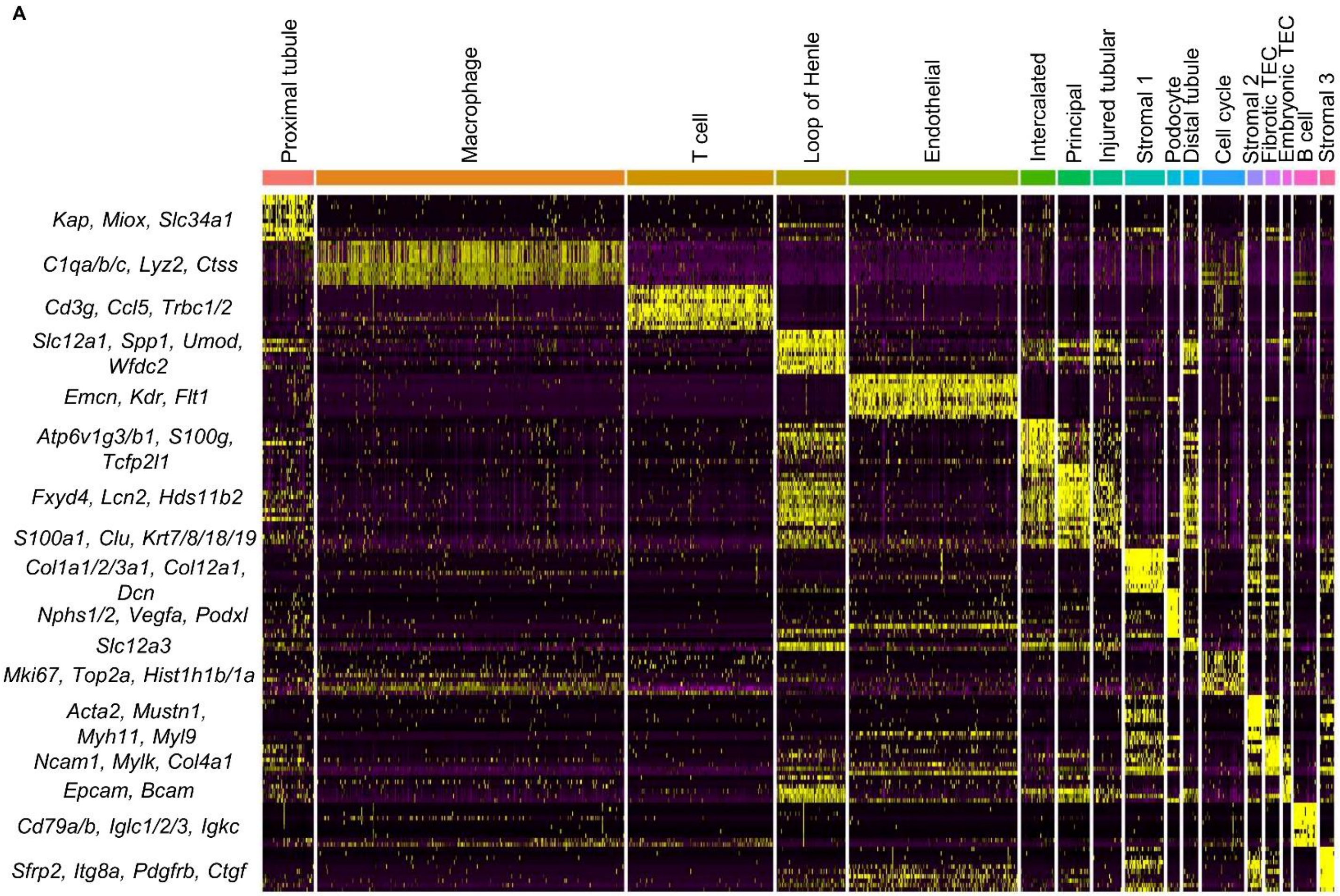

**Supplemental figure 6. Invasive hemodynamic reveals no significant systemic cardiovascular function changes caused by UIR and UUO compared to control. (A)** Heart rate in beats per minute (bpm) at baseline, with vasoconstrictive (PE, phenylephrine), vasodilative (SNP, sodium nitroprusside) and inotropic agent (DOB, dobutamine). Ctrl – control interval between agents. Data are presented as mean values ± SD, n=3-4 per group. **(B)** Mean arterial pressure (mmHg) at baseline, with vasoconstrictive (PE, phenylephrine), vasodilative (SNP, sodium nitroprusside) and inotropic agent (DOB, dobutamine). Ctrl – control interval between agents. Data are presented as mean values ± SD, n=3-4 per group. **(C)** Systolic pressure (mmHg) at baseline, with vasoconstrictive (PE, phenylephrine), vasodilative (SNP, sodium nitroprusside) and inotropic agent (DOB, dobutamine). Ctrl – control interval between agents. Data are presented as mean values ± SD, n=3-4 per group.

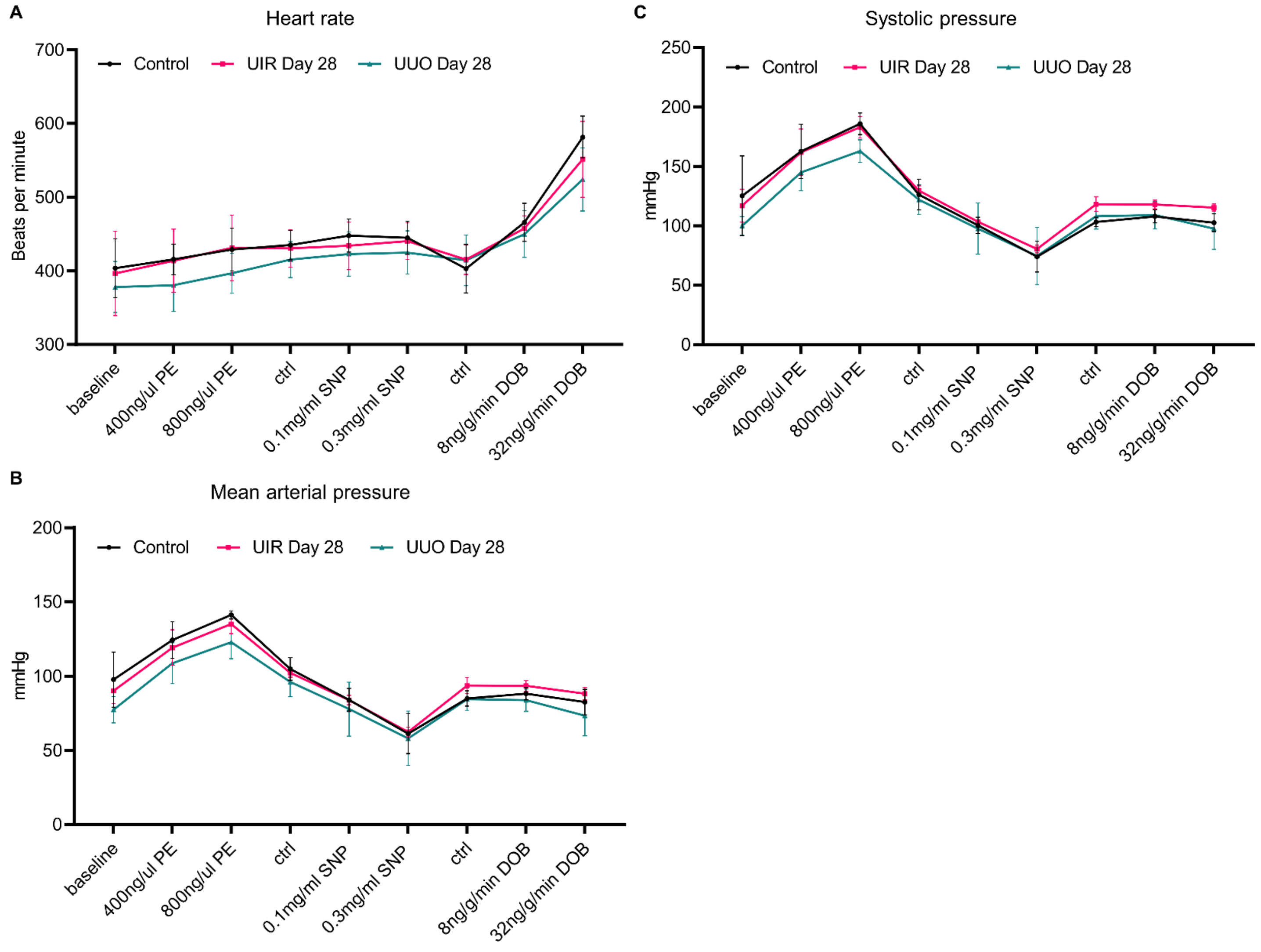

**Supplemental Figure 7. Drop-seq reveals molecular and cellular landscapes of kidney fibrosis in an independent cohort of UIR and UUO mice.** (A) Feature plots demonstrating tubular, endothelial, podocyte, immune and three distinctive stromal clusters in UIR and UUO models of kidney fibrosis. **(B)** GO Biological process of “Stromal 1, 2 and 3” marker genes vs other populations in UIR, – log2 (pValue). **(C)** GO Biological process of “Stromal 1, 2 and 3” marker genes vs other populations in UUO, – log2 (pValue).

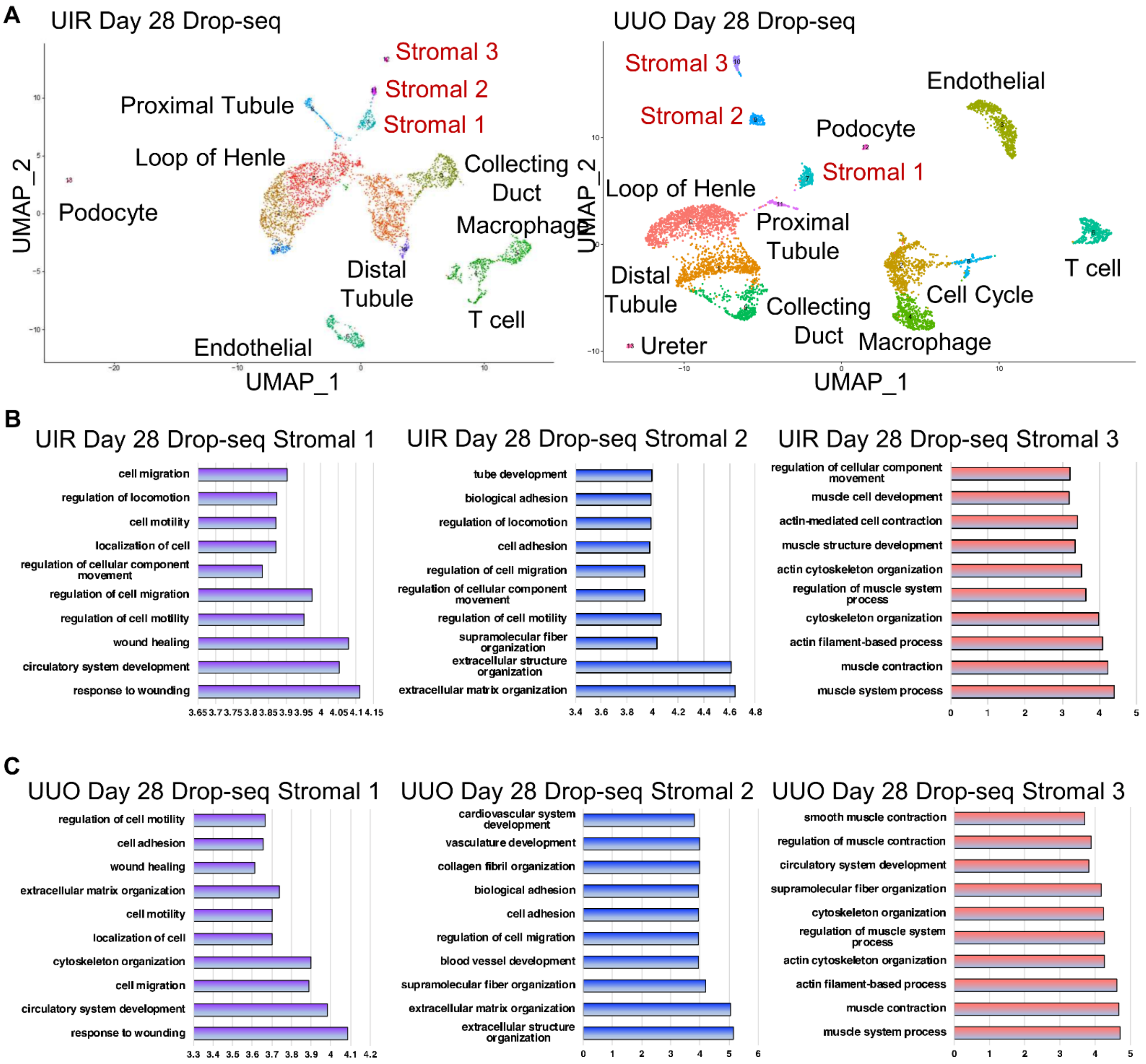

**Supplemental Figure 8. Doublet removal does not affect scRNA-seq identified cellular landscapes in the normal and fibrotic kidneys.** (A) UMAP shows renal cell populations in the control, UIR and UUO kidneys (n=3-5 per group) after doublet removal. Clusters are distinguished by different colors. PT, proximal tubules, LOH, loop of Henle, DT, distal tubule, CD-P, collecting duct principal, CD-I, collecting duct intercalated, Podo, podocytes, Endo, endothelial, Macro, macrophages, Str, stromal, TECs, tubular epithelial cells, E-TEC, embryonic tubular epithelial cells.

A

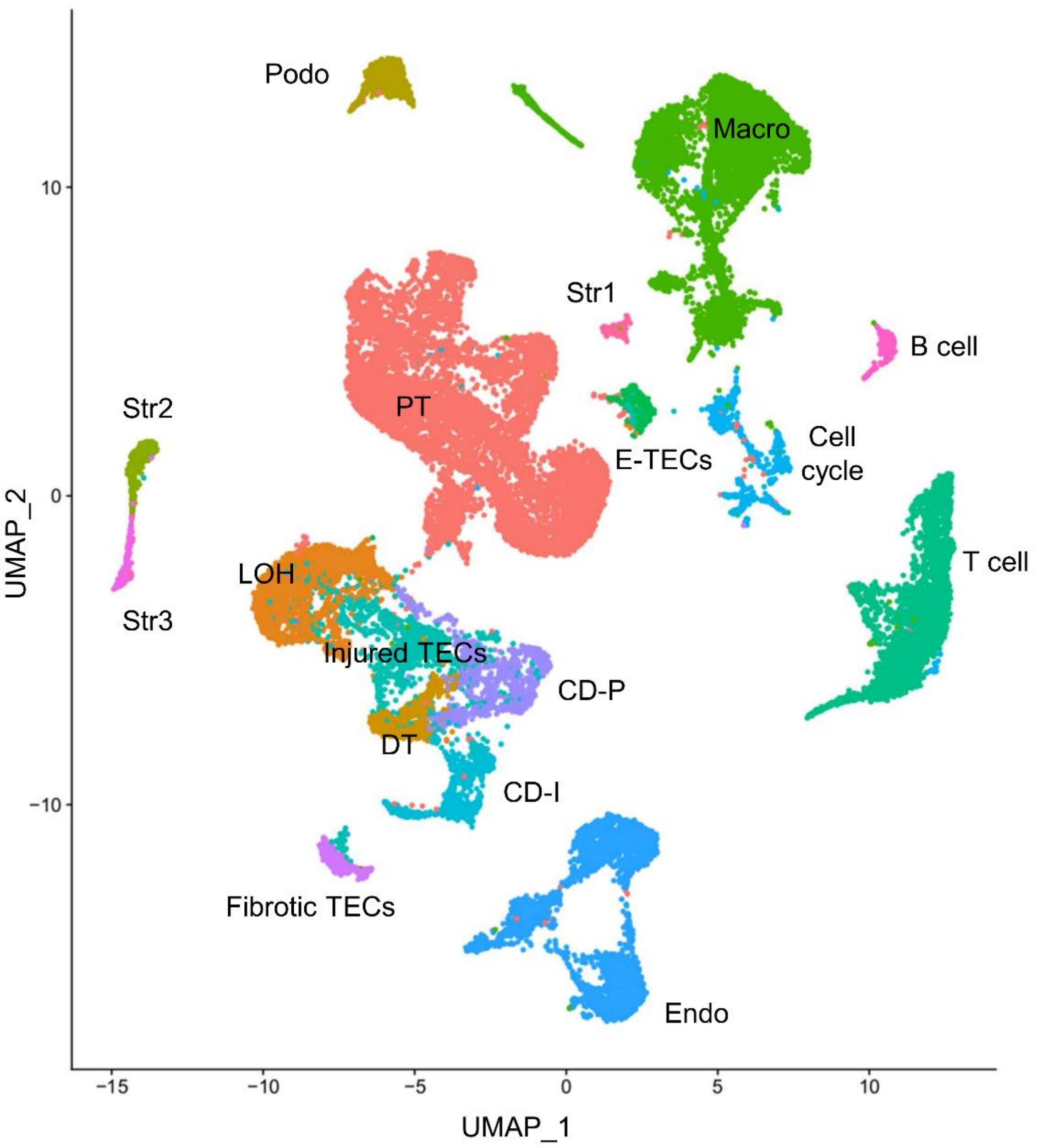

**Supplemental figure 9. scRNA-seq reveals transcriptional identity of fibrotic TECs.**

**(A)** Feature plots of *Cdh16* expression in the tubular clusters, including UIR and UUO fibrotic TECs. Gene expression levels are color-coded. **(B)** Feature plots of epithelial developmental genes *Osr2* and *Cdh6* expression in fibrotic and embryonic TECs. Gene expression levels are color-coded.

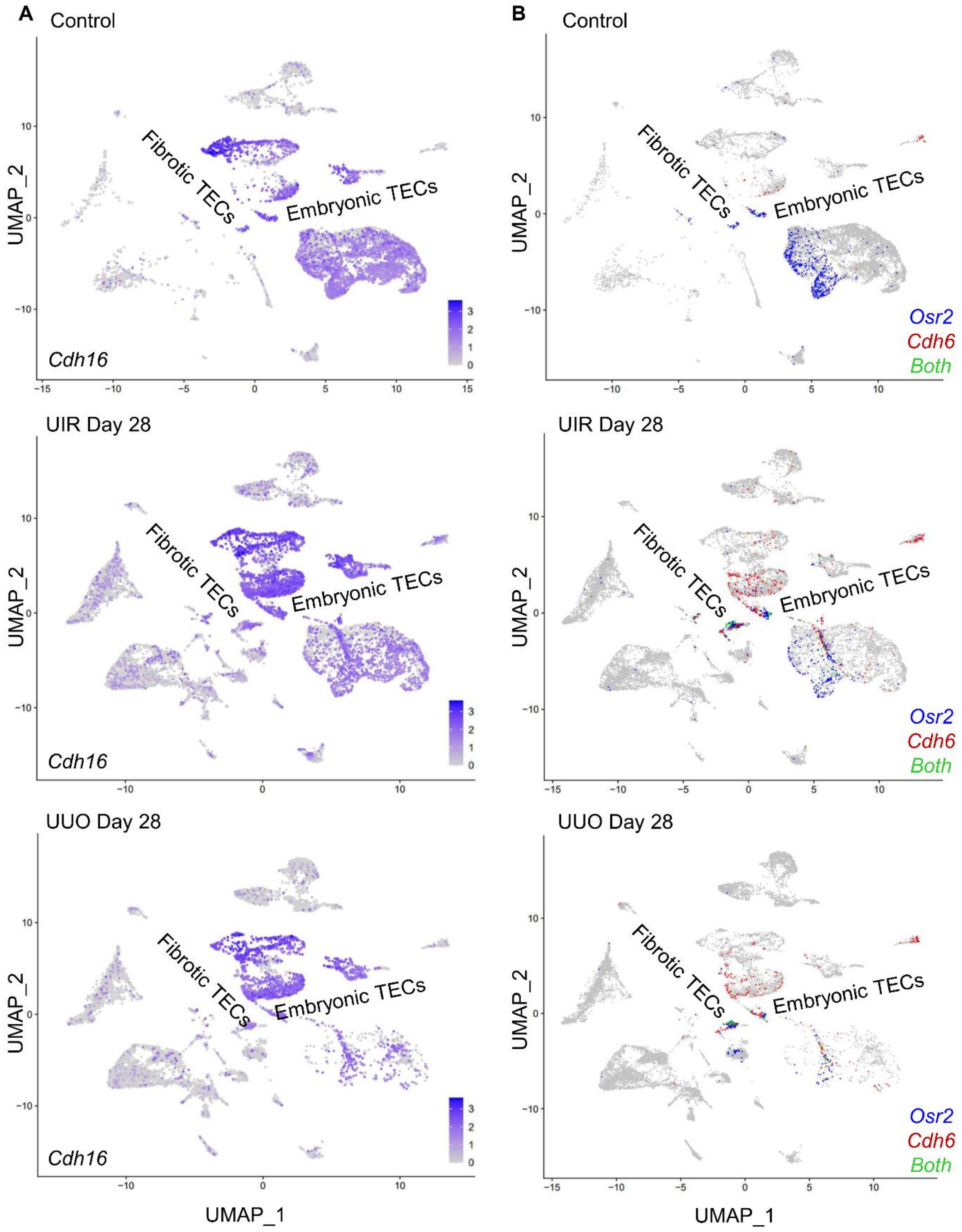

**Supplemental figure 10. Integrated trajectory analysis shows little evidence of EMT in advanced renal injury. (A)** Integrated trajectory analysis of tubular (proximal tubule, loop of Henle, embryonic and fibrotic TECs) and stromal clusters of control, UIR and UUO Day 28. Number 1 represents significant branch point of differentiation. **(B)** Integrated trajectory analysis of the control, UIR and UUO stromal, endothelial and fibrotic TEC populations. Number 1 represents significant branch point of differentiation.

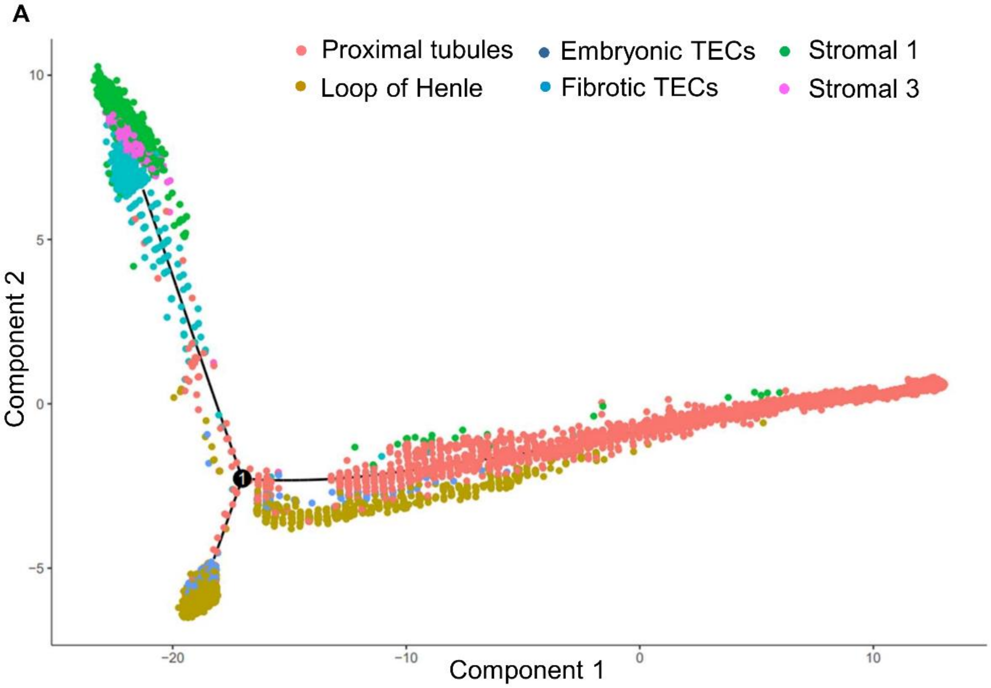


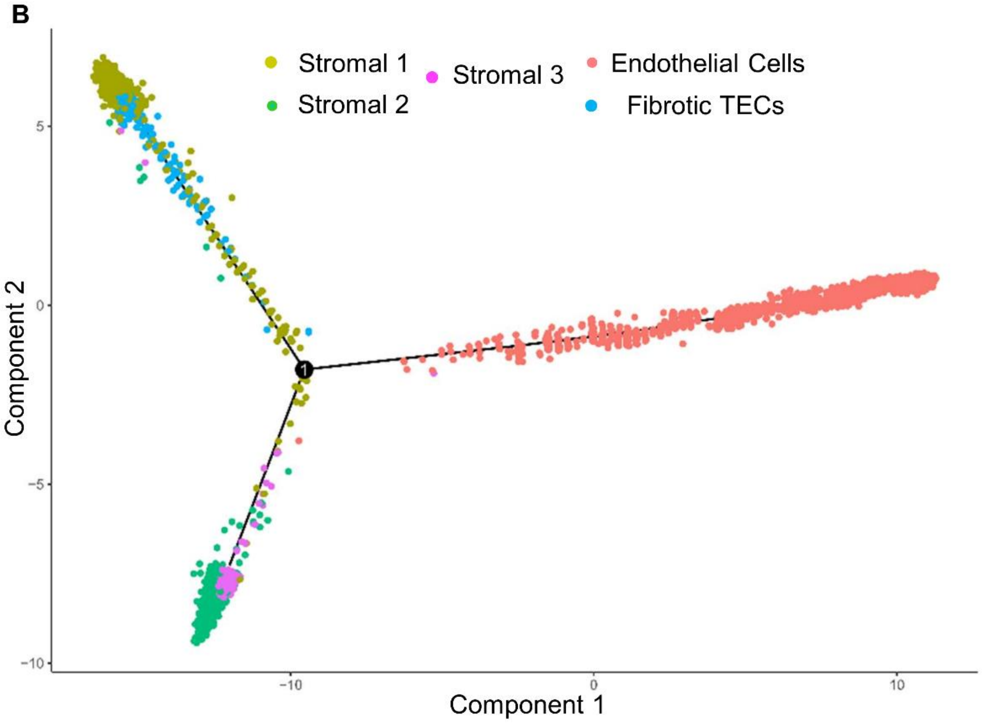

**Supplemental figure 11. scRNA-seq integrated trajectory analysis reveals transcriptional identities of kidney ECM producing stromal and tubular clusters.**

**(A)** The linear heatmap shows gene expression changes in the Control, UIR and UUO Day 28 stromal and fibrotic TEC clusters over a Pseudo – Time. The cells are clustered based upon similarity of gene expression. Gene expression level is color coded.

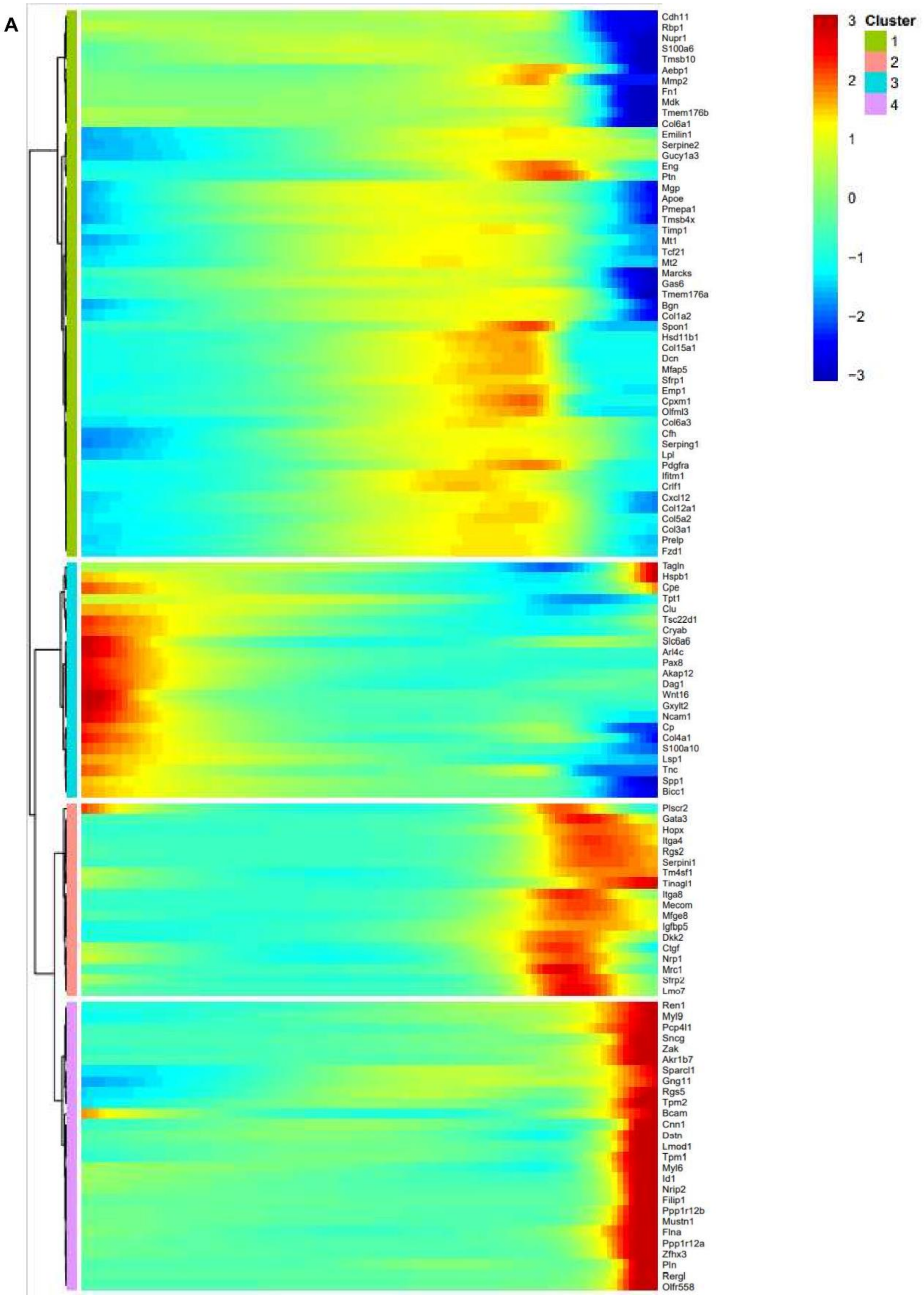

**Supplemental figure 12. scRNA-seq demonstrates stromal markers *Pdgfrb* and *Gli1* expression in normal and fibrotic kidneys. (A)** and **(B)** Feature plots of *Pdgfrβ* and *Gli1* in stromal and fibrotic TEC populations of control, UIR and UUO kidneys. Gene expression levels are color-coded. Str, stromal, Podo, podocyte.

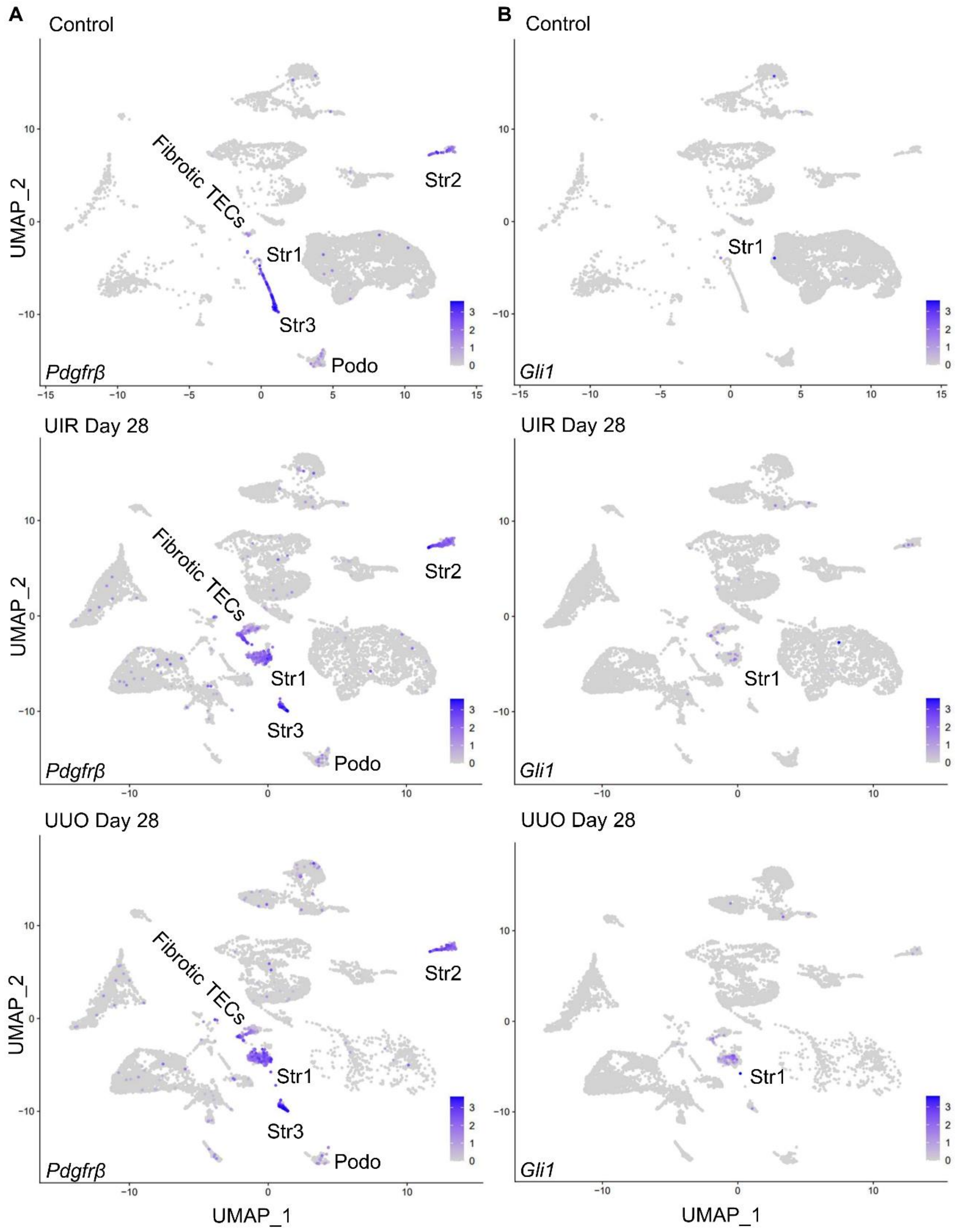

**Supplemental figure 13. Gucy1a3 specifically labels activated kidney fibroblasts in the control and fibrotic kidney scRNA-seq datasets after doublet removal. (A)** Feature plots demonstrate *Gucy1a3, Acta2* and *Col1a1* expression in the control and fibrotic kidneys after doublet removal. Note that *Acta2* and *Col1a1* are elevated in the tubular, immune and endothelial compartments, including pro-fibrotic TECs, while *Gucy1a3* expression remains restricted to three activated fibroblast clusters with negligent tubular expression. *Gucy1a3* expression levels are color-coded.

A

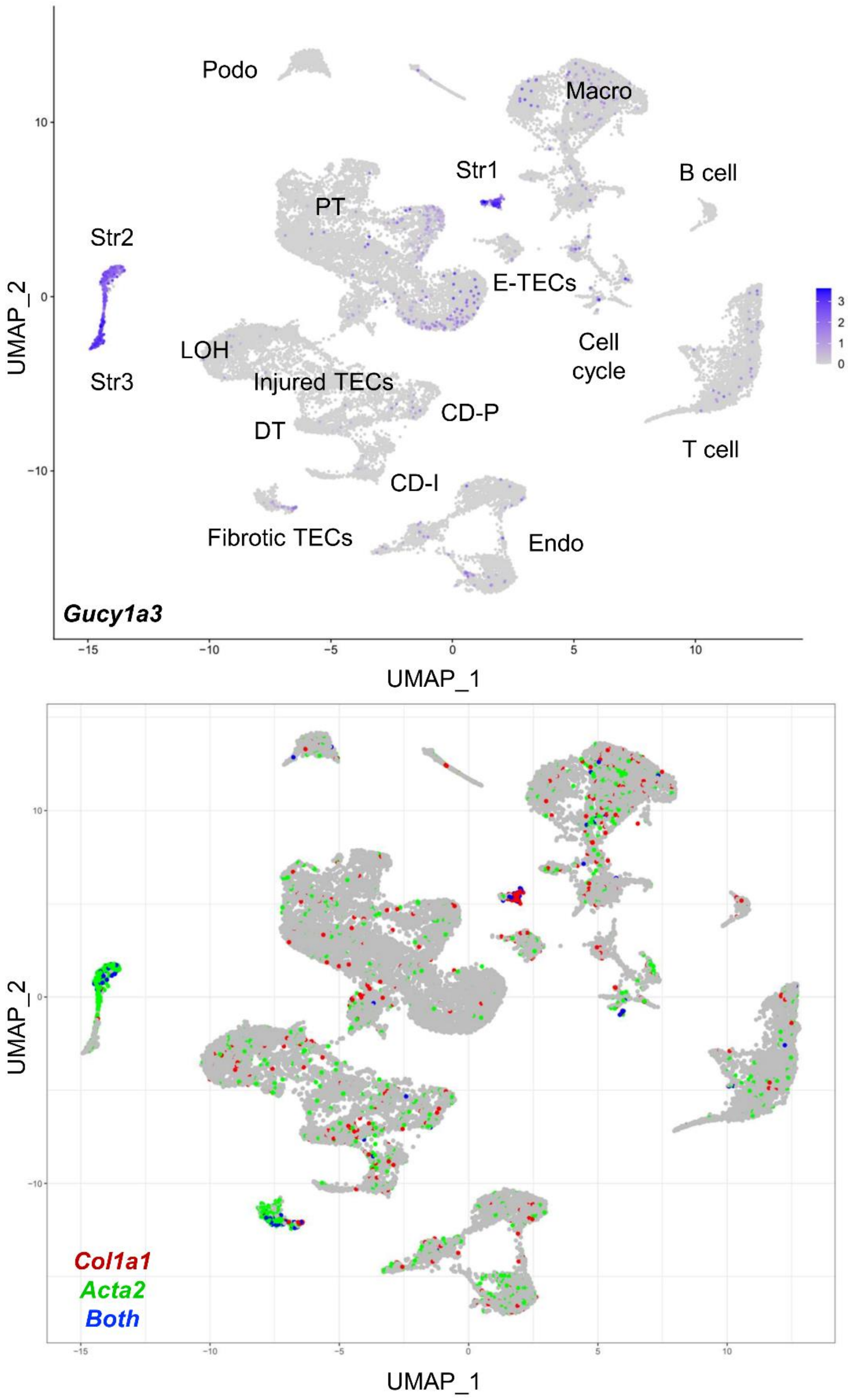

**Supplemental figure 14. scRNA-seq dissects the novel cellular and molecular mechanisms of crosstalk between the stromal and epithelial cells in the UIR model of kidney fibrosis.**

(A) Circos plot of ligand-receptor interactions between proximal tubules (salmon), fibrotic TECs (heliotrope), embryonic TECs (magenta), stromal 1 (iris blue), stromal 2 (light slate blue) and stromal 3 (rose) clusters in control kidney. The populations producing putative ligand are shown at blunt end of the arrow; the populations producing putative receptor are highlighted with the color code line next to the ligand-producing cluster. Arrow points from the ligand-producing to receptor-producing populations. The names of all putative ligands and receptors with respect to the cell populations are available in Supplementary data 4. (B) Circos plot of ligand-receptor interactions between proximal tubules (salmon), fibrotic TECs (heliotrope), embryonic TECs (magenta), stromal 1 (iris blue), stromal 2 (light slate blue) and stromal 3 (rose) clusters in UIR kidney. (C) qPCR of *Col18a1* expression in control and UIR kidneys, n=6 per group. (D) Representative images and quantification of endostatin (*Col18a1* C-terminal product) Western blot, control and UIR kidneys, n=4-5 per group. (E) Representative images of Col18a1 IHC in control and UIR kidneys. Upper images – original magnification, ×10, 0.64 μm/px zoom; lower images - original magnification, ×40, 0.16 μm/px zoom into the upper images, area highlighted with black frames. *pValue ± 0.05, ****pValue ≤ 0.0001 compared to control, Student's *t* test, (C) and (D).

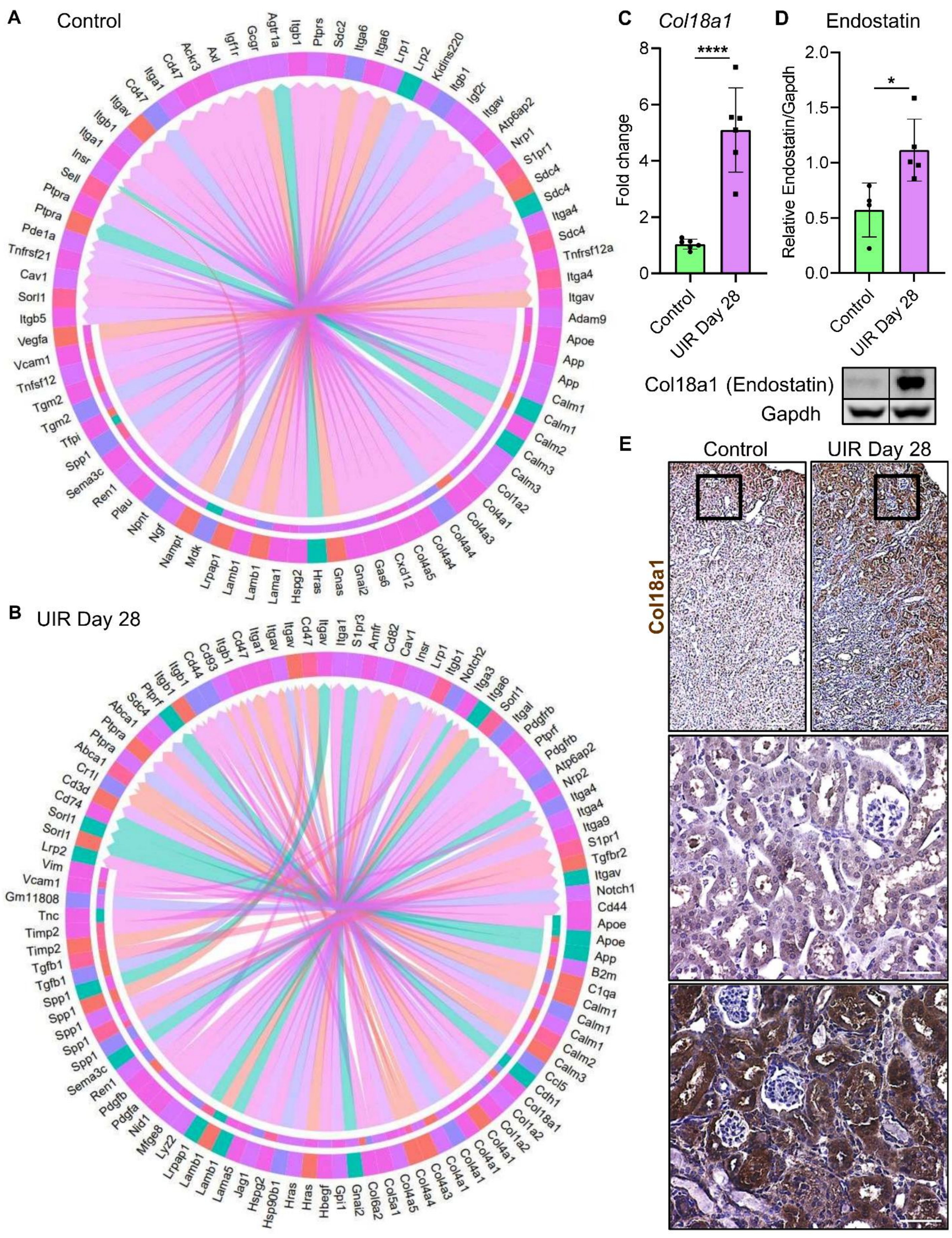
A
Control
B
UIR Day 28
Agtr1a
Itgb1
Ptprs
Sdc2
Itga6
Lrp1
Lrp2
Kidins220
Igf2r
Itgav
Atp6ap2
Nrp1
S1pr1
Sdc4
Itga4
Tnfrsf12a
Adam9
Apoe
App
Calm1
Calm2
Calm3
Col1a2
Col4a1
Col4a3
Col4a4
Col4a5
Cxcl12
Gas6
Gnai2
Gnas
Hras
Hspg2
Lama1
Lamb1
Lrpap1
Mdk
Nampt
Ngf
Npnt
Plau
Ren1
Sema3c
Spp1
Tfpi
Tgm2
Tnfsf12
Vcam1
Vegfa
Itgb5
Sorl1
Cav1
Tnfrsf21
Pde1a
Ptpra
Sell
Insr
Itga1
Itgav
Cd47
Ackr3
Axl
Igf1r
Gcgr
Cd44
Cd93
Amfr
Cd82
Notch2
Itga3
Itgal
Pdgfrb
Ptprf
Nrp2
Itga9
Tgfbr2
Notch1
B2m
C1qa
Ccl5
Cdh1
Col18a1
Col5a1
Col6a2
Gpi1
Hbegf
Hsp90b1
Jag1
Lama5
Lyz2
Mfge8
Nid1
Pdgfa
Pdgfb
Tgfb1
Timp2
Tnc
Gm11808
Vim
Cd74
Cd3d
Cr1l
Abca1
C
Col18a1
Fold change
****
Control
UIR Day 28
D
Endostatin
Relative Endostatin/Gapdh
*
Control
UIR Day 28
Col18a1 (Endostatin)
Gapdh
E
Control
UIR Day 28
Col18a1

**Supplemental figure 15. scRNA-seq demonstrates *Col18a1* expression in normal and fibrotic kidneys.**

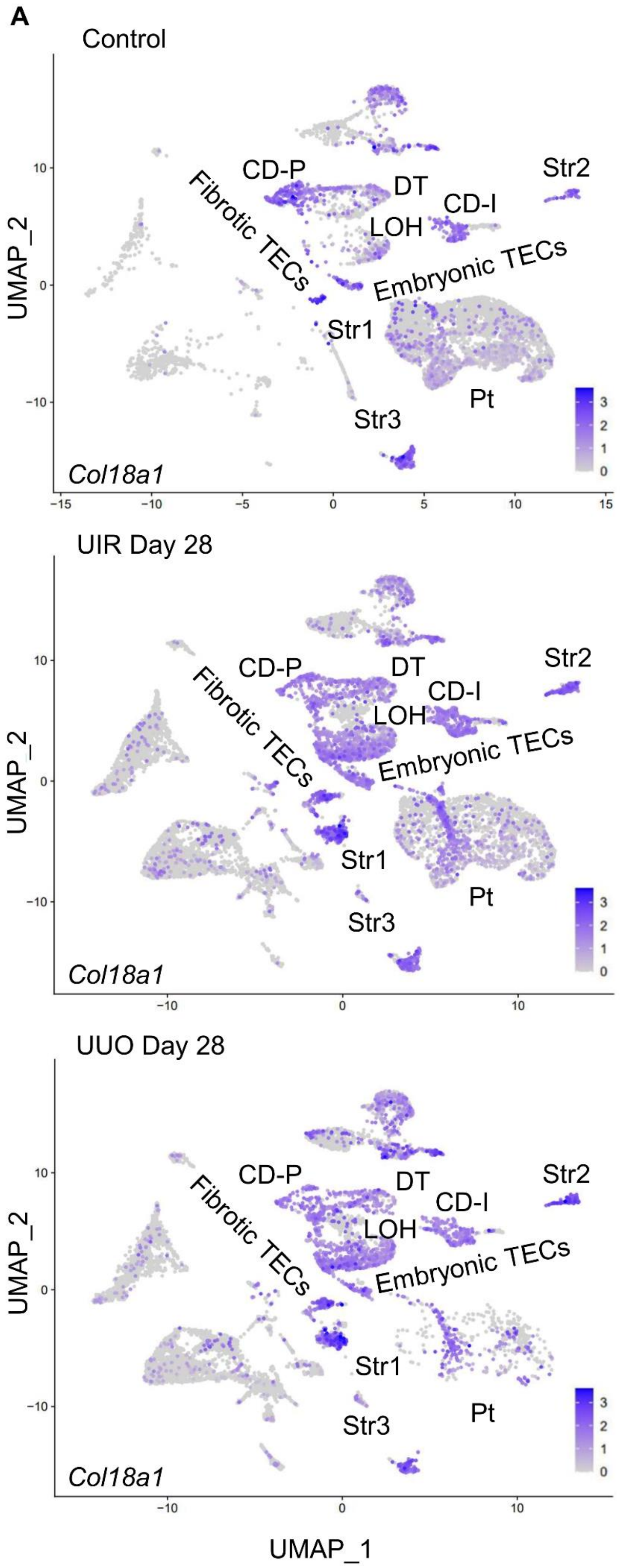


**(A)** Feature plots of *Col18a1* expression in stromal and tubular populations of control, UIR and UUO kidneys, including fibrotic and embryonic TECs. Gene expression levels are color-coded. PT, proximal tubules, LOH, loop of Henle, DT, distal tubule, CD-P, collecting duct principal, CD-I, collecting duct intercalated, Str, stromal, TEC, tubular epithelial cells.

**Supplemental figure 16. scRNA-seq reveals novel gene expression signatures in tubular epithelial cells in kidney fibrosis.** (A) ToppCluster analysis of biological processes enriched among the genes elevated in proximal tubules of UIR and UUO over the control. Genes related to “Collagen binding” are highlighted with purple polygons, “ECM binding” – tyle, both – dark blue. Genes related to “Cadherin binding”, including Ahnak (shown in red), are highlighted with green polygons, “Cell adhesion molecule binding” – pink, “Structural molecule activity” – light blue. Complete lists of genes are presented in Supplementary data 5. (B) ToppCluster analysis of biological processes enriched among the genes elevated in fibrotic TECs of UIR and UUO over the control. Genes related to “Collagen binding” are highlighted with purple polygons, “ECM binding” – tyle, “ECM structural constituent” – pink. Genes related to “Cadherin binding”, including Ahnak (shown in red), are highlighted with blue polygons, “Cell adhesion molecule binding” – green, “Structural molecule activity” – peach. (C) Representative images of combined *Spp1* CISH (black) and Krt8 IF (red) in control, UIR and UUO kidneys. Mild stromal Krt8 expression in normal kidney is highlighted with green arrows. *Spp1* elevation in UIR and UUO Krt8-positive injured tubules is shown with yellow arrows. Original magnification, ×40, 0.17 μm/px zoom. Complete lists of genes are presented in Supplementary data 5, (A) and (B).

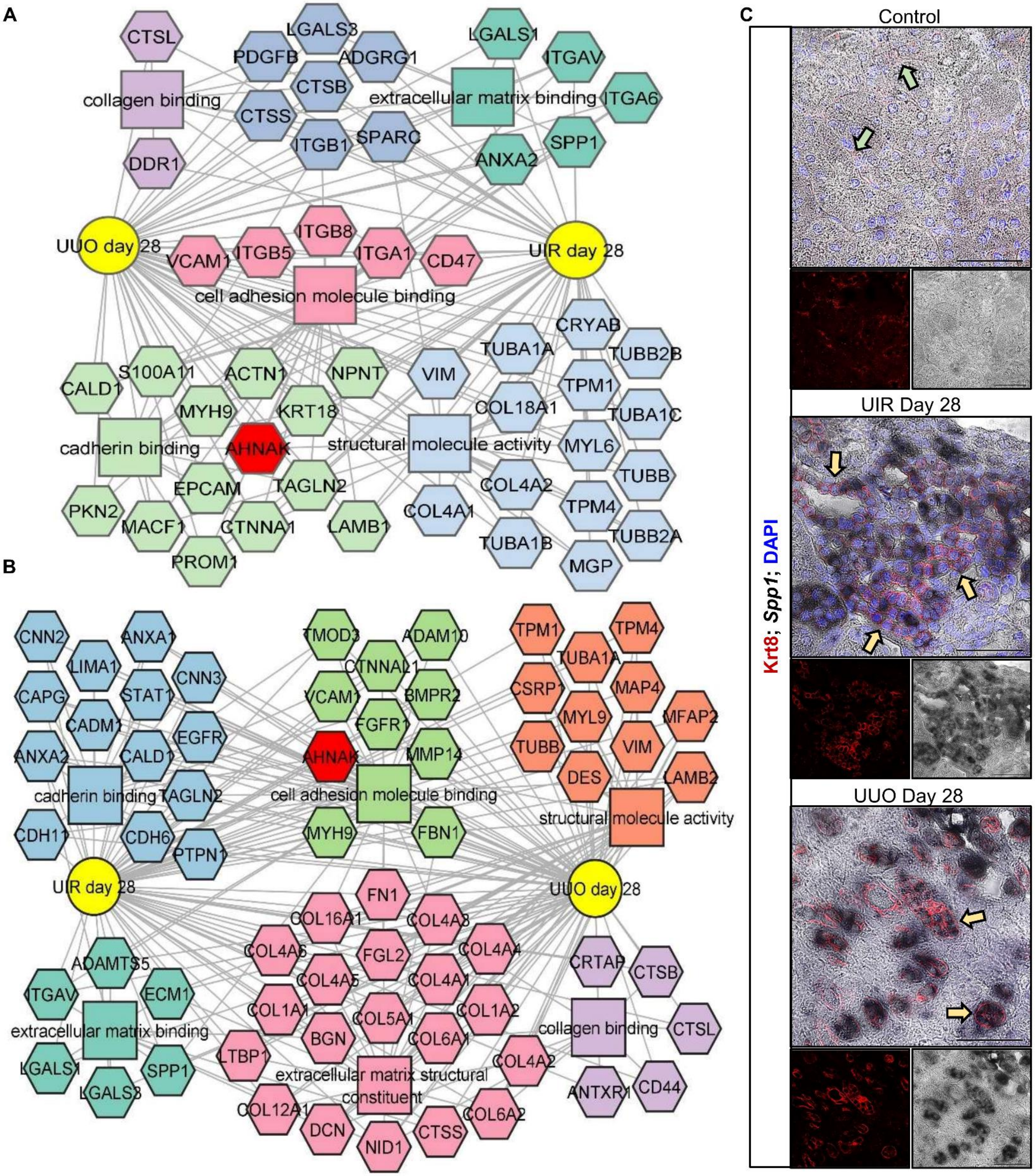
A
CTSL
collagen binding
DDR1
LGALS3
PDGFB
ADGRG1
CTSB
CTSS
ITGB1
SPARC
LGALS1
ITGAV
extracellular matrix binding
ITGA6
SPP1
ANXA2
UUO day 28
UIR day 28
ITGB8
VCAM1
ITGB5
ITGA1
CD47
cell adhesion molecule binding
CRYAB
TUBA1A
TUBB2B
S100A11
ACTN1
NPNT
VIM
TPM1
CALD1
MYH9
KRT18
COL18A1
TUBA1C
cadherin binding
AHNAK
structural molecule activity
MYL6
EPCAM
TAGLN2
COL4A2
TUBB
PKN2
MACF1
CTNNA1
LAMB1
COL4A1
TPM4
TUBA1B
TUBB2A
PROM1
MGP
B
CNN2
ANXA1
LIMA1
CNN3
CAPG
STAT1
CADM1
EGFR
ANXA2
CALD1
cadherin binding
TAGLN2
CDH11
CDH6
PTPN1
TMOD3
ADAM10
CTNNAL1
VCAM1
BMPR2
FGFR1
AHNAK
MMP14
cell adhesion molecule binding
MYH9
FBN1
TPM1
TPM4
TUBA1A
CSRP1
MAP4
MYL9
MFAP2
TUBB
VIM
DES
LAMB2
structural molecule activity
UIR day 28
UUO day 28
FN1
COL16A1
COL4A3
COL4A6
FGL2
COL4A4
COL4A5
COL4A1
COL1A1
COL5A1
COL1A2
BGN
COL6A1
LTBP1
extracellular matrix structural constituent
COL4A2
COL12A1
DCN
NID1
CTSS
COL6A2
ADAMTS5
ITGAV
ECM1
extracellular matrix binding
LGALS1
SPP1
LGALS3
CRTAP
CTSB
collagen binding
CTSL
ANTXR1
CD44
C
Control
UIR Day 28
UUO Day 28
Krt8; Spp1; DAPI

**Supplemental figure 17. CKD models elicit enduring elevation of AKI-inducible genes *Myh9* and *Sh3bgrl3*. (A)** and **(B)** Feature plots of *Myh9* and *Sh3bgrl3* expression in control, UIR and UUO kidney cell populations. Gene expression levels are color-coded. FT, fibrotic TECs, ET, embryonic TECs, PT, proximal tubules, LOH, loop of Henle, DT, distal tubule, CD-P, collecting duct principal, CD-I, collecting duct intercalated, Str, stromal, TEC, tubular epithelial cells. **(C)** RT-qPCR of *Myh9* and *Sh3bgrl3* in control, UIR and UUO kidneys, n=4-6 per group, Student's *t* test, ***pValue ≤ 0.001, ****pValue ≤ 0.0001 compared to control.

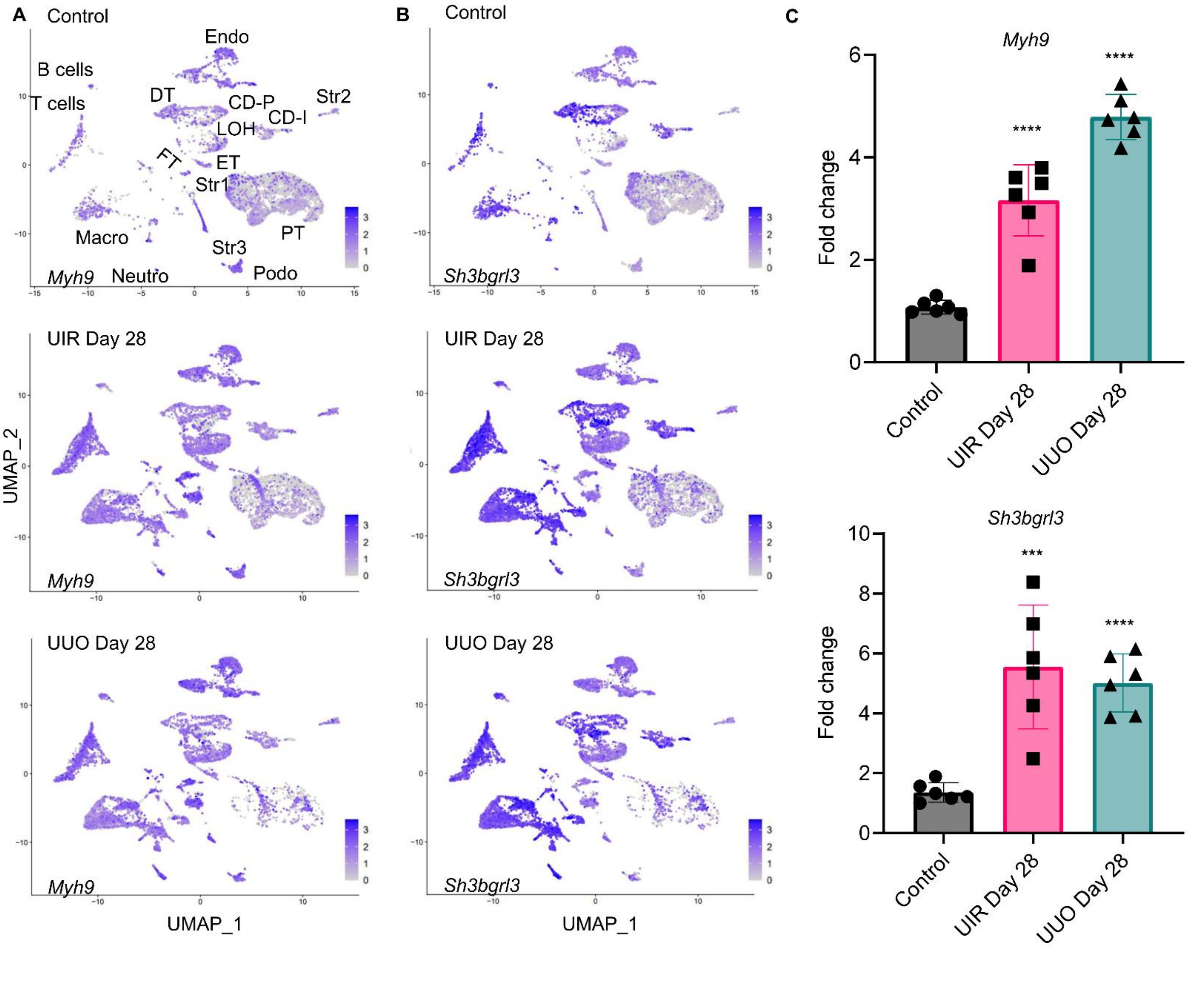

**Supplemental figure 18. Western blot validates UIR and UUO inducible Ahnak elevation in fibrotic kidneys. (A)** Original image of Western blot which demonstrates Ahnak elevation in both UIR and UUO Day 28 kidneys. Ahnak was detected with Proteintech 16637-1-AP antibody which we tested using siRNA induced AHNAK knockout in primary human renal proximal tubular epithelial cells (RPTECs). Bands selected as representative are shown with yellow arrows.

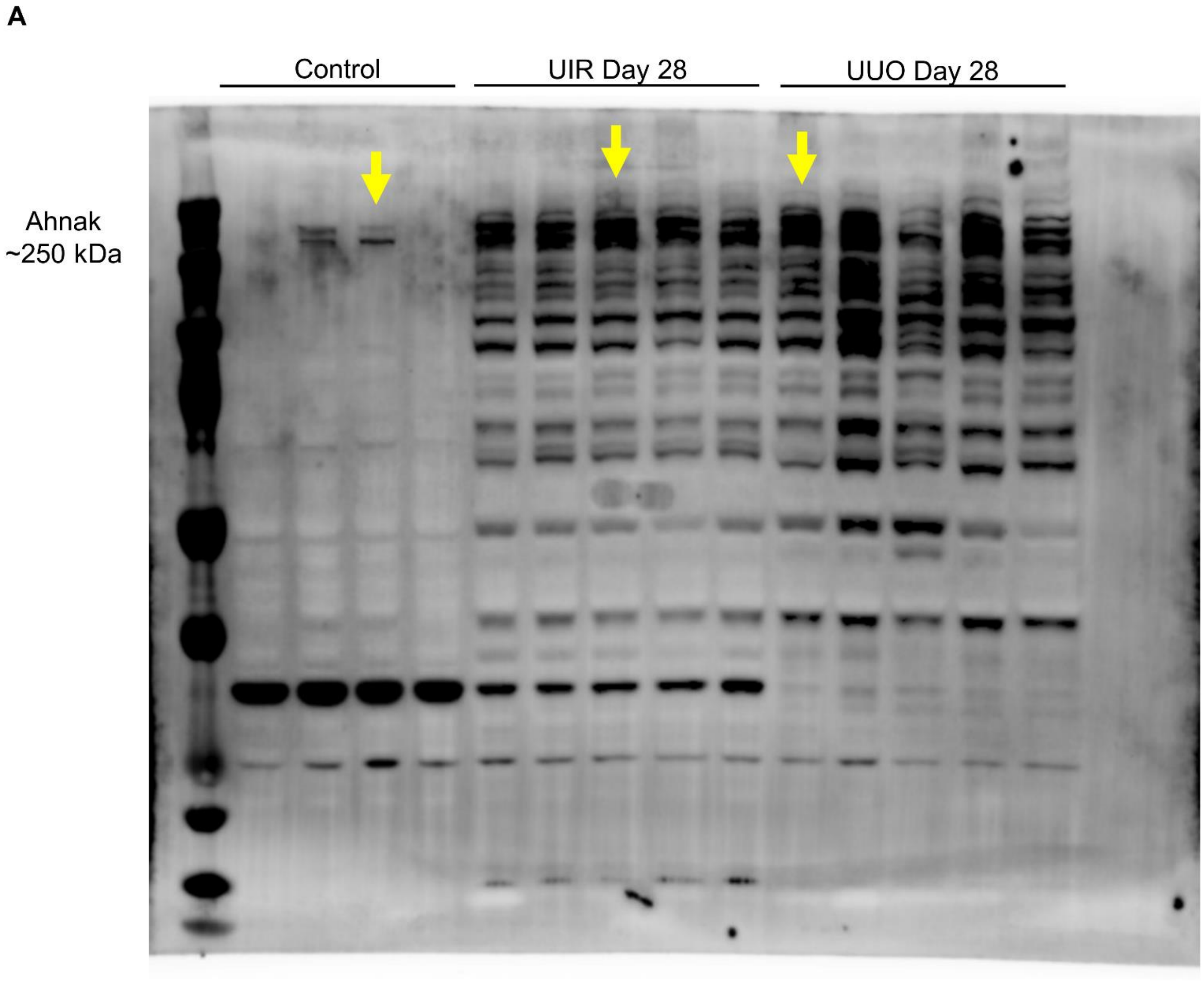

**Supplemental figure 19. Western blot validates AHNAK siRNA knockdown. (A)** Original image of Western blot which demonstrates AHNAK expression in RPTECs cultured on the baseline, with vehiche (Vh), with non-targeting siRNA-Scramble or targeting siRNA-AHNAK constructs, with or without TGFβ. Ahnak was detected with Proteintech 16637-1-AP antibody (1 to 250 dilution for 48 hours). Bands selected as representative are shown with black arrows. **(B)** Original image of Western blot which demonstrates equal housekeeping protein expression (GAPDH) in RPTECs cultured on the baseline, with vehiche (Vh), with non-targeting siRNA-Scramble or targeting siRNA-AHNAK constructs, with or without TGFβ.

A

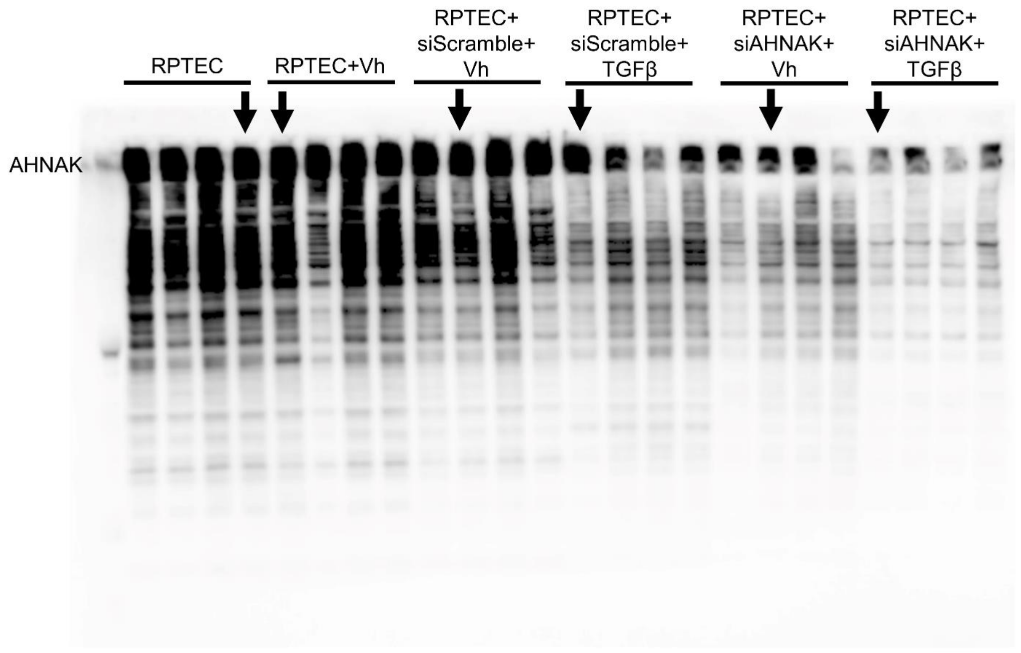


B

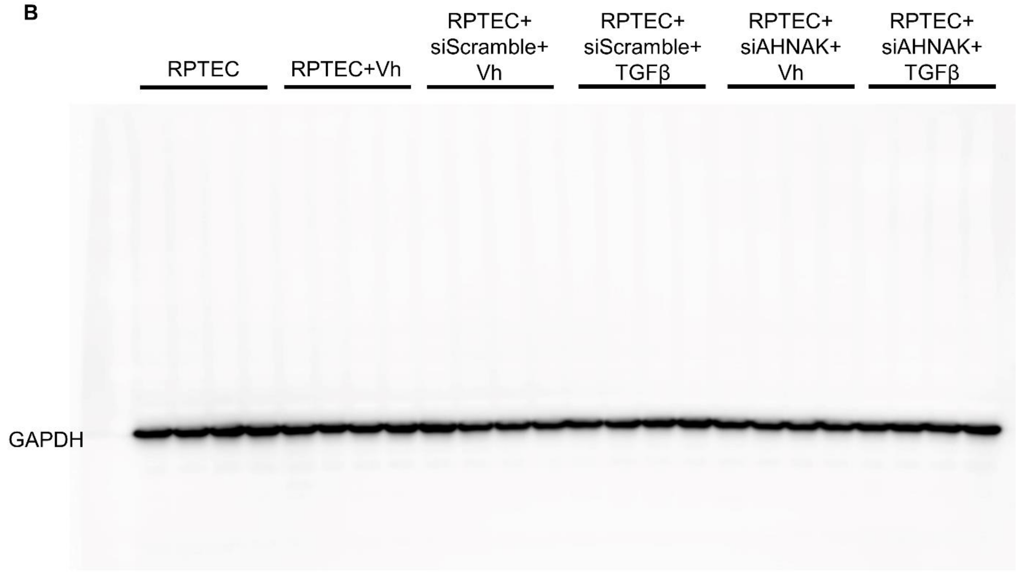
